# Hummingbirds rapidly respond to the removal of vision and control a sequence of rate-commanded maneuvers in milliseconds

**DOI:** 10.1101/2023.12.13.571331

**Authors:** Md Zafar Anwar, Bret W. Tobalske, Suyash Agrawal, Jean-Michel Mongeau, Haoxiang Luo, Bo Cheng

## Abstract

Hummingbird flight is the epitome of extreme aerial agility and controlled stability, as hummingbirds routinely exercise a variety of stunning aerobatic feats. Yet, the control of these amazing maneuvers is not well understood. Here we examined how hummingbirds control a sequence of maneuvers within milliseconds and tested whether and when their vision is active during this rapid process. We elicited escape flight in calliope hummingbirds and removed visible light at various instants during the maneuvers and quantified their flight kinematics and responses. We show that the escape maneuvers were composed of rapidly-controlled sequential modules, including evasion, reorientation, nose-down dive, forward flight, and nose-up to hover. The hummingbirds did not respond to the light removal during evasion and reorientation until a critical light-removal time; afterward they showed two categories of luminance-based responses that rapidly altered maneuvering modules to terminate the escape. We also show that hummingbird maneuvers are rate-commanded and require no active braking, i.e., their body angular velocities were proportional to the change of wing motion patterns, a trait that likely alleviates the computational demand on flight control. Together, this work uncovers the key traits of hummingbird agility, which can also inform and inspire designs for next-generation agile aerial systems.

## Introduction

The ability to fly is a hallmark of evolutionary success in nature and technological breakthroughs by humans (1). Both nature and engineering have explored various forms of aerodynamic forces (2–8), not merely to defy gravity, but also to achieve aerial locomotion with combined stability and agility for a wide range of speeds. Among all natural and human-made fliers, hummingbird flight represents an extreme form of high agility and controlled stability at low flight speed (9). These tiny vertebrate fliers routinely exercise a variety of aerobatic feats with swiftness, precision, and astonishing control, which are put into full display when they fight to defend territories (10), dance in courtship display (11), or escape from threats (12, 13). A casual observer can easily distinguish hummingbirds from other fliers by these unique aerobatic feats.

Hummingbirds excel not only in individual aerobatic maneuvers but also in their ability to rapidly generate and combine a sequence of maneuvers (9), for example, during escape flight (12– 14) and competitive interactions (15). To achieve such complex aerobatic feats, the birds need both the muscle capacity (16) to generate accelerations that quickly change body orientation and speed and the flight control (14) that guarantees stability within individual maneuvers and swiftness in transitions among maneuvers. Presumably, this ability emerges from a complex interaction between their flight control system (17) (including neural, sensory, and muscular processes) and physical system (18, 19) (including aerodynamics, body, and wing dynamics). Previous studies have shown that hummingbirds possess elevated muscle capacity and large force vectoring ability for maneuvering (14, 16); however, little is known about flight control and the role of vision within a maneuvering sequence, and how they rapidly modulate wing motion patterns and body movement.

This study is motivated to improve our understanding of hummingbird agility by providing quantitative results regarding flight control and the role of vision during rapid aerobatic maneuvers. We focused on escape flight, which is an ecologically relevant behavior including a sequence of rapid maneuvers with near-maximal performance (12, 13). First, we elicited escape flight repeatedly in calliope hummingbirds flying in a flight chamber (see Materials and Methods). In the middle of each escape flight, we impeded the birds’ vision by turning off visible ambient LED light at various electronically-controlled time instants and recorded the birds’ entire maneuvering sequence using highspeed cameras under infrared light (Figure 1A). Using this method, we were able to test whether the vision was active during various phases of the maneuvers, how its removal altered the maneuvers, and whether and how hummingbirds stabilized in the dark.

**Figure 1.**
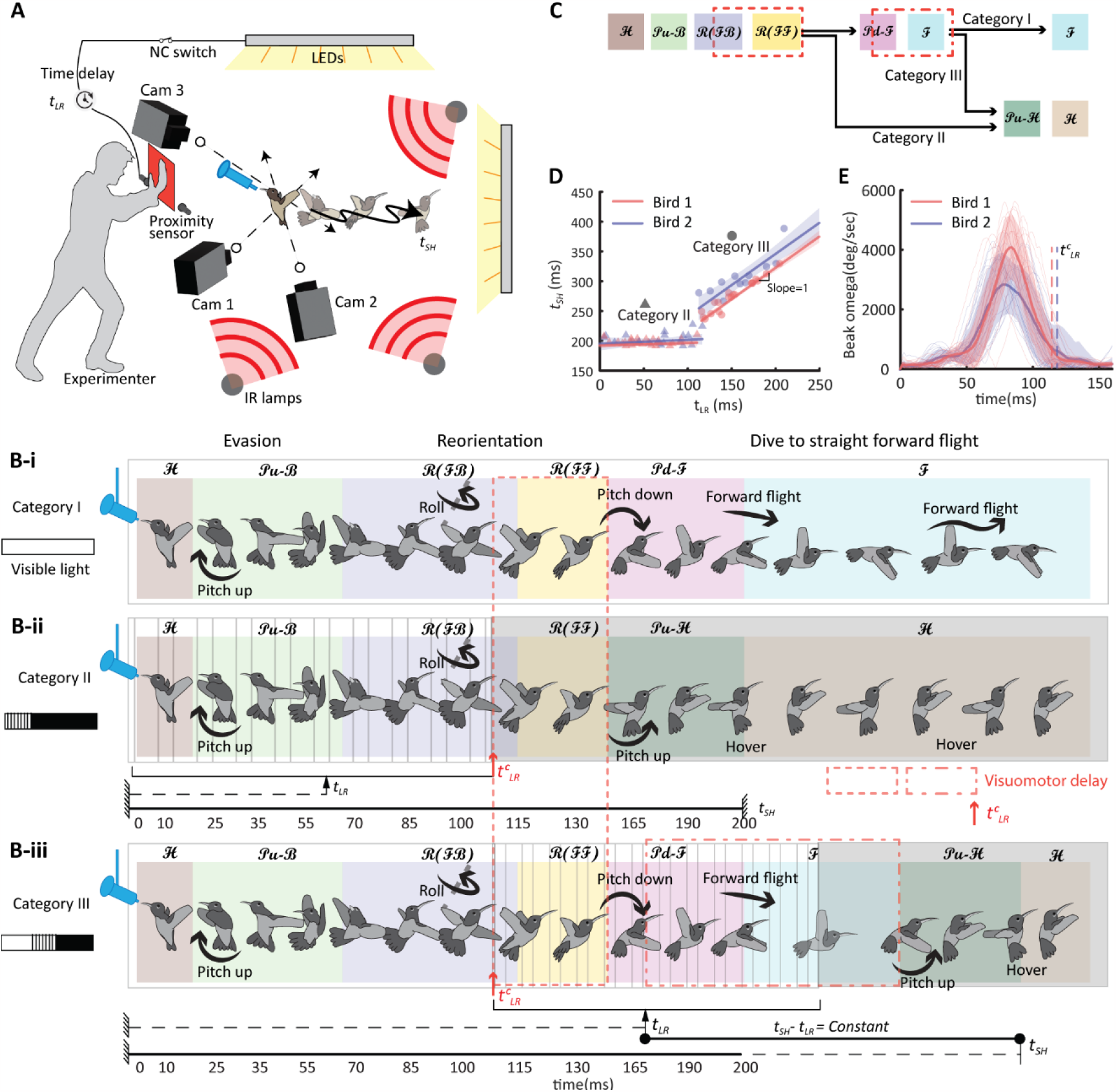
Escape maneuvering modules and luminance-based responses in calliope hummingbirds. (A) Experimental setup. A hummingbird was hovering and feeding at a feeder in a long corridor, and it was startled by an experimenter using a red board to elicit escape flight. A proximity sensor was used to detect the movement of the red board, based on which it turned off the power to visible LED lights at varied time delays using a nominally-closed switch, thereby removing the visible light of hummingbirds at different time instants of the escape flight. The entire flight sequence was recorded using three high-speed monochrome cameras under infrared lights. (B) Three categories of escape behaviors according to luminance-based responses. The grated region indicates the time window when the visible light was removed at a variable time instant *t*_*LR*_. The grey region indicates the time window when the visible light was always absent to birds. Note that any solid horizontal line represents a fixed time window, and any dashed horizontal line represents a variable time instant. The red dashed boxes represent the time window from visual input to initiation of 𝓅***𝓊-ℋ***, which represents the visuomotor delay. The colored background patches indicate the maneuvering modules. (B-i) Category I: visible light was never removed (indicated by the white box). (B-ii) Category II: - visible light was removed within the grated box [0 **–** 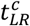] at variable *t*_*RL*_. (B-iii) Category III: visible light was removed at a variable time 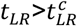 (grated region). Note that in this category, the bird always used a constant amount of time (*t*_*SH*_-*t*_*LR*_) from the light removal instant to hover in the dark, while the duration of forward flight ***ℱ*** is variable depending on *t*_*LR*_. (C) Categories of escape maneuvers and the corresponding sequences of maneuvering module according to the luminance-based responses. Maneuvering modules include hover (***ℋ***), evasive maneuver (𝓅***𝓊-ℛ***), reorientation (***ℛ***), nose down dive (𝓅***𝒹-ℱ***), nose up the pitch (𝓅***𝓊-ℋ***), and forward flight (***ℱ***). The red dashed boxes are the same as that shown in the previous sub-figure. (D) Relationship between light removal time instant (*t*_*LR*_) and the time instant to stop and hover (*t*_*SH*_). (E) Instantaneous head saccade rate during the escape flight. The dashed vertical line shows the critical light removal instance,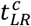.

Next, we identified maneuvering modules of the escape flight sequence and showed how they were sequenced according to the luminance-based response to the light removal at various timings. We then analyzed the three-dimensional wing kinematics of the entire maneuvering sequence, from which we extracted Wing Motion Primitives (WMPs)(20) and their temporal combinations with varying magnitudes (see Materials and Methods). We evaluated how the combination of WMPs generate body rotations using cross-correlation between instantaneous WMP magnitude and the body angular rates. In addition, we assessed whether and when the wing kinematics followed a predetermined, stereotyped pattern, by calculating the statistical similarity among wing kinematics from different escape flights (see Materials and Methods). Using this method, we provided predictions on the varying degree of feedforward control (high similarity) vs feedback control (low similarity). Collectively, our results show that hummingbirds have active vision and can rapidly change their body and wing motion during highly-dynamic maneuvers (e.g., up to 6 times in 250ms). The angular rates of hummingbird maneuvers can be directly commanded by the changes of wing motion patterns (which we referred to as *rate-commanded maneuvers*, a term borrowed from the literature on aircraft flight (21, 22)), this reveals the simplicity in hummingbird flight control.

## Results

### Maneuvering modules of escape flight without light removal (category I)

We analyzed a total of 22 trials for bird 1 and 21 trials for bird 2 (see *SI Appendix*). We observed three categories of escape flight (Figure 1B and Movies S1-5), including category I for escape trials without light removal, and category II and III for two categories of the birds’ responses to light removal at variable time instants (*t*_*LR*_). In category I (Figure 1B-i and Movie S3), where visible light was present all time, the maneuvers were composed of four maneuvering modules: an initial evasive maneuver (***𝒫𝓊-ℬ***, 2-4 wingbeats, with body ***𝒫***itch ***𝓊***p and ***ℬ***ackward acceleration), followed by a fast reorientation (***ℛ***, 5-9 wingbeats, with rapid (***ℛ***oll rotation), then a dive (***𝒫𝒹-ℱ***, 10-12 wingbeats, with ***𝒫***itch ***𝒹***own and ***ℱ***orward acceleration,) into forward flight (***ℱ***). The evasive maneuver (***𝒫𝓊-ℬ***) allowed the bird to quickly exit the original hovering location (peak backward acceleration: 11.9 ± 3.6 m/s^2^, bird 1 and 11.8 ± 2.3 m/s^2^, bird 2) and redirect its body’s long axis towards the direction of escape (peak pitch rates: 1294 ± 390°/s, bird 1 and 1691 ± 283°/s, bird 2; peak yaw rate: 879 ± 297°/s, bird 1, and 970 ± 320°/s, bird 2; also see *SI Appendix*, Figure S7). The reorientation (***ℛ***) included a rapid head saccade followed by a body roll (peak roll rate: 2541 ± 512°/s, bird 1, and 2609 ± 361°/s, bird 2; also see *SI Appendix*, Figure S7) and allowed the birds to completely turn away from the threat while regaining a stable body posture. Finally, the pitch-down dive and forward flight (***𝒫𝒹-ℱ)*** enabled the bird to attain high speed to fly away from the threat.

### Luminance-based responses that altered maneuvering modules (category II and III)

Once the escape was initiated, the birds did not respond to the light removal applied during the evasion (***𝒫𝓊-ℬ***) and reorientation (***ℛ***), and they completed these modules in darkness without losing stability (the patterns, initiation and termination timings of these two maneuvering modules were not affected by the light removal timing or the light conditions) (Figure 1B). However, the subsequent forward dive (***𝒫𝒹-ℱ***) exhibited variable initiation and termination timings depending on the luminance-based responses to continue or terminate the escape.

In category II (Figure 1B-ii and Movie S4), when the visible light was removed within a critical light-removal time from the initiation of escape 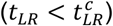, ***𝒫𝒹-ℱ*** was not initiated (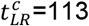 ms<u>^1^</u> for bird 1 and 116 ± 3 ms for bird 2) (see Materials and Methods). Instead, the birds terminated the escape as they ***𝒫***itched ***𝓊***p and returned to ***ℋ***over (***𝒫𝓊-ℋ***) in the dark. The time that the birds came to a complete stop and hover (i.e., stop-hover time *t*_*SH*_) is nearly fixed (*t*_*SH*_ = 193 ± 5 ms for bird 1 and 199 ± 10 ms for bird 2). Notably, 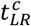 occurred near the end of the head saccade when the birds’ gaze was stable (Figure 1E), concurrent with the end of the reorientation module.

In category III (Figure 1B-iii and Movie S5), when 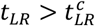, the birds continued to escape as the ***𝒫𝒹-ℱ*** was initiated. However, the ***ℱ*** was eventually terminated after a nearly constant visuomotor delay from the instants of light removal (*t*_*LR*_) (78 ± 9ms for bird 1 and 91 ± 20ms for bird 2); and then the birds performed ***𝒫𝓊-ℋ*** to hover in the dark at *t*_*SH*_ (*t*_*SH*_ *− t*_*LR*_= 119 ± 7 ms for bird 1 and 143 ± 17 for bird 2). The duration of the forward flight, therefore, varied linearly with the *t*_*LR*_ (Figure 1D and Materials and Methods).

The above results show that the birds generated two categories of luminance-based, binary responses that altered the sequence of maneuvering modules, depending on the *t*_*LR*_ (Figure 1C).

The first response was whether to initiate ***𝒫𝒹-ℱ*** to continue escape (categories I and III, with illumination at 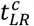) or ***𝒫𝓊-ℋ*** to terminate escape (category II without illumination at 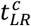). It can be shown that the first response was dependent on the luminance only at 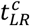, or whether visual input was present or removed at 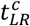, independent of the specific light removal instance 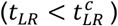 (see *SI Appendix*). The total visuomotor delay from 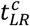 to the initiation of ***𝒫𝓊-ℋ*** or ***𝒫𝒹-ℱ*** was 39 ± 3 ms (bird 1) and 30 ± 4 ms (bird 2), which includes the time needed for visual processing (and potentially decision-making), motor control, and wing musculoskeletal response. The second response was whether to continue ***ℱ*** and escape (category I) or initiate ***𝒫𝓊-ℋ*** to terminate escape (category III). This response was generated with a total visuomotor delay of 78 ± 9ms (bird 1) and 91 ± 20ms (bird 2) from the light removal timing (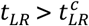, note the birds’ gaze was stable in this period). In addition, these results also show that hummingbirds can continue to maneuver, stabilize, and hover in the dark.

It is not clear why the hummingbird did not respond to the light removal during the evasion and reorientation prior to 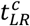. There are two possibilities for this observation: either the bird can’t alter its trajectory of evasion and reorientation once the escape is initiated (visually open loop), or the bird can alter the trajectory but in all the demonstrations actively decided not to. Notably, 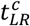occurring at the end of the head saccade suggests that the bird can only respond to the light removal with a stable gaze, however, our results also show that the birds didn’t respond to the light removal prior to the head saccade after the initiation of the escape maneuver (e.g., between 20-50ms, Figure 1E). Therefore, it remains inconclusive whether the unresponsiveness of the birds during evasion and reorientation can be attributed to unstable gaze.

### Wing motion primitives for evasion and reorientation

To initiate the escape, the birds rapidly switched wing motion patterns (Figure 2B-i) to produce hover (***ℋ***) → evasion (***𝒫𝓊-ℬ***) → reorientation (***ℛ***) maneuvering sequence in less than 150 ms. We extracted the wing motion primitives (WMPs)(20) for these maneuvering modules and estimated how they were combined wingbeat-by-wingbeat during the maneuvers (see Materials and Methods). We first defined hover Wing Motion Primitive (WMP^***ℋ***^) based on averaging the wingbeat motion patterns preceding the evasion. To obtain the maneuvering WMPs during the ***ℋ*** → ***𝒫𝓊-ℬ*** → ***ℛ***, we time-aligned the trajectories from all the analyzed demonstrations (N=22 for bird 1, and N=21 for bird 2) and estimated a single intended trajectory (23) for each bird (see *SI Appendix*). We then measured the statistical difference (Hellinger distance (24)) between the WMP^***ℋ***^ and each maneuvering wingbeat from the estimated intended trajectory; and we defined the WMPs for evasion (WMP^***𝒫𝓊-ℬ***^) and reorientation (WMP^***ℛ***^) at the local maxima (WMP^***𝒫𝓊-ℬ***^ at 3^rd^ wingbeat and WMP^***ℛ***^ at 8^th^ wingbeat, Figure 2B). Next, assuming an arbitrary wingbeat pattern during the escape maneuver is a linear combination of WMP^***ℋ***^, WMP^***𝒫𝓊-ℬ***,^ and WMP^***ℛ***^, we calculated the wingbeat-by-wingbeat magnitude of each WMP (20) (Figure 3A-ii). Assuming the birds’ wing musculoskeletal system effected motor control into changes of wing motion within a wingbeat (25, 26), we hypothesized that the amplitude of each WMP reflected the magnitude of whole-muscle and motor-unit recruitment for the corresponding WMP (i.e., a measure of the collective intensity of muscle activation).

**Figure 2.**
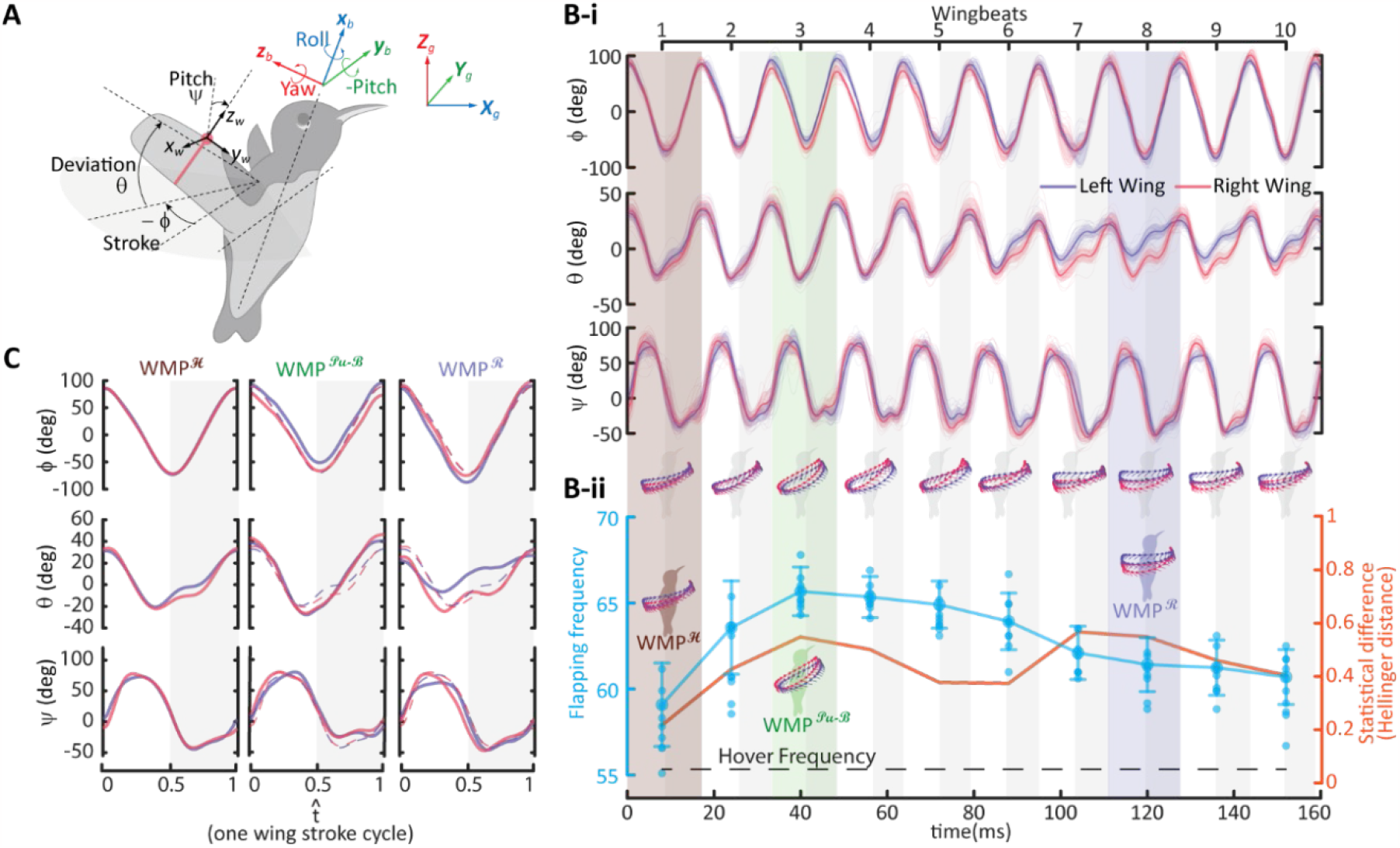
Wing kinematics and Wing Motion Primitives (WMPs) during the escape maneuvers. (bird 1, see bird 2 result in *SI Appendix*, Figures S9-10). (A) Illustration of the body coordinate frame *{x*_*b*_, *y*_*b*_, *z*_*b*_*}* and wing coordinate frame *{x*_*w*_, *y*_*w*_, *z*_*w*_*}*. The wing kinematics are defined by angles stroke(*ϕ*), deviation(*θ*), and wing pitch(*ψ*). (B-i) Wing kinematics (intended trajectory) during ***ℋ*** → 𝓅***𝓊-ℬ*** → ***ℬ*** maneuvering modules. The shaded areas enclosing the curves indicate ±1 s.d. (22 trials). (B-ii) Wing flapping frequency during escape flight (cyan) and individual wingbeat probabilistic distance from hovering wing kinematics (orange), which measures the statistical difference between individual maneuvering wingbeat and averaged hovering wingbeat. The brown, green, and purple patches represent the wingbeats at which WMP^***ℋ***^, WMP^***𝓅𝓊-ℬ***^, and WMP^***ℛ***^are defined, respectively. (C) Wing kinematic profile for WMPs, dashed curves in WMP^***𝓅𝓊-ℬ***,^ and WMP^***ℛ***^represent the hovering kinematics (WMP^***ℋ***^) for comparison. The horizontal axis represents the normalized time for a single wing stroke cycle. Downstroke periods are indicated by gray-shaded regions.

**Figure 3.**
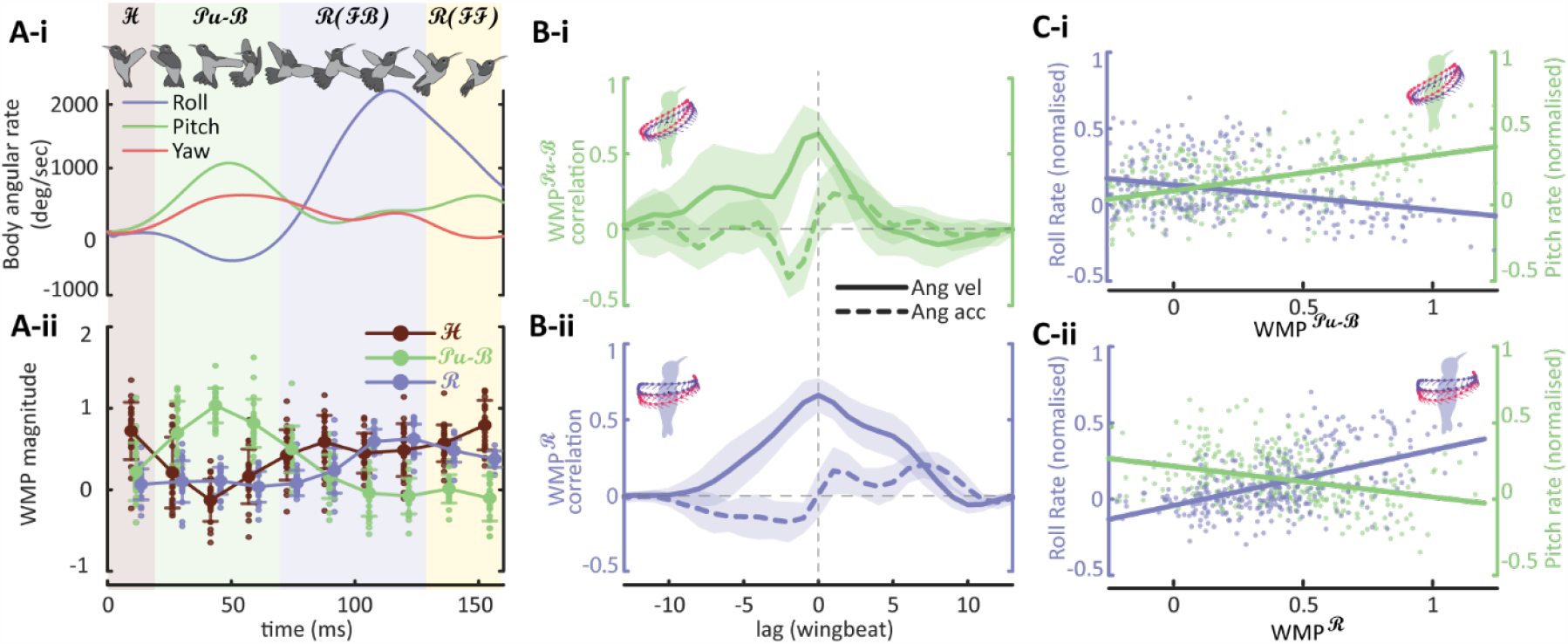
Wing Motion Primitives (WMPs) are phase-locked with body angular rate. (bird 1, see bird 2 result in *SI Appendix*, Figure S11). (A-i) Body angular rate in body fixed frame. (A-ii) WMP magnitude of individual wingbeats during escape flight. The WMP magnitude follows the trend of the respective body angular rate, i.e., WMP^***𝓅𝓊-ℬ***^ follows the pitching rate while WMP^***ℛ***^ follows the roll rate. Background patches denote the maneuvering modules as referenced in Figure 1. (B) The cross correlation between WMPs magnitude and respective cycle-averaged angular velocity and acceleration. It is the angular velocity, instead of angular acceleration, that correlates strongly with 0 wingbeat lag and is phase-locked with WMPs. (C) Relationship between WMPs magnitude and respective normalized angular rate, and their linear regression. The slopes of linear fit are significantly different from 0. The pitch rate relates proportionally to WMP^***𝓅𝓊-ℬ***^ and inversely to WMP^***ℛ***^, while the roll rate relates proportionally to WMP^***ℛ***^and inversely to WMP^***𝓅𝓊-ℬ***^.

### Body rotations rate-commanded by wing motion primitives

We then examined how the combinations of the three WMPs produced ***ℋ*** → ***𝒫𝓊-ℬ*** → ***ℛ*** body maneuvering sequence, a process that involves highly unsteady aerodynamics (27) and nonlinear body dynamics (28, 29). The results showed that the magnitude of WMP^***𝒫𝓊-ℬ***^ was nearly phase-locked with the cycle-averaged body pitch rate (rather than pitch acceleration), throughout the entire maneuvering sequence. Similarly, the reorientation WMP^***ℛ***^ was nearly phase-locked with the cycle-averaged body roll rates (Figure 3A). Moreover, cross-correlation analyses showed that it was the body angular rate, instead of angular acceleration, that correlated strongly to WMPs with zero wingbeat lag (Figure 3B). In addition, the WMPs never reversed polarity when the birds decelerated from body rotation (Figure 3A), indicating that there was no active braking. These results showed that hummingbirds’ body angular rates responded, within a wingbeat cycle, proportionally to the change of wing motions (or WMP magnitudes), which showed that their angular maneuvers are rate-commanded(22) by wing motion patterns. A detailed discussion distinguishing rate-commanded and non-rate-commanded maneuvers can be found in *SI Appendix*.

### Cross-demonstration similarity of wing motion patterns during *ℋ* → *𝒫𝓊-ℬ* → *ℛ*

We calculated the temporal variation of the cross-demonstration similarity of wing motion patterns throughout maneuvering modules ***ℋ*** → ***𝒫𝓊-ℬ*** → ***ℛ*** (Figure 4A), as a way to predict the degree of feedforward vs feedback control of wing motion (see discussion). The similarity index was calculated using the Coefficient of Multiple Correlation (CMC) (30, 31) within a sliding window of two wingbeats (Equation 3) (Materials and Methods), sliding with an increment of half wingbeat. We also calculated the variance of body orientation (measured as the angle between body long axis and gravitational axis, Figure 4B) to examine its potential relationship with the wing motion.

**Figure 4.**
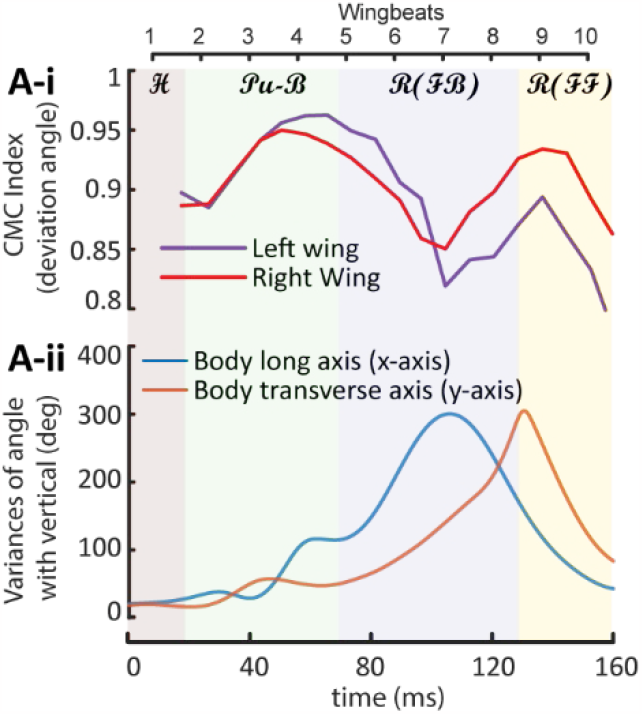
Wing kinematics similarity (CMC) indicates the degree of feedforward and feedback flight control during escape maneuvers. (bird 1, see bird 2 result in *SI Appendix*, Figure S12). (A-i) Coefficient of Multiple Correlation (CMC) for measuring the similarity amongst instantaneous wing kinematics of different escape demonstrations. Color background patches denote the maneuvering modules as referenced in Figure 1. (A-ii) Variances in the body orientation during escape maneuvers. The cyan curve denotes the variance in the angle between the bird’s body’s long axis (x-axis) and the global vertical axis, and the orange curve denotes the variance in the angle between the bird’s body transverse axis (y-axis) and the global vertical axis.

**Figure 5.**
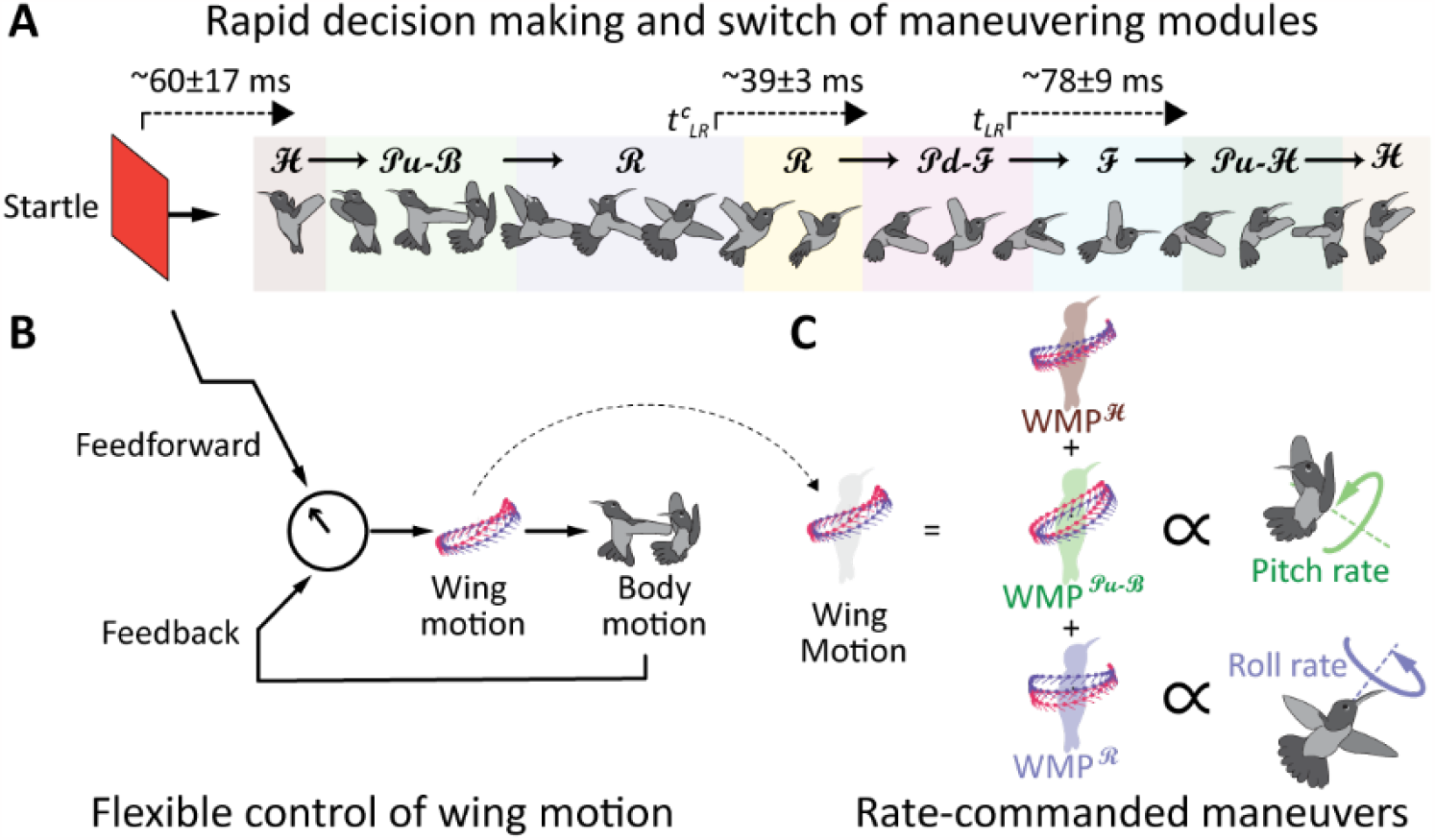
Summary of key traits of hummingbird agility. A) Birds were able to switch the maneuvering modules as many as 6 times in less than 250ms with low latency in the visuomotor mapping. B) Bird’s body angular rate was rate-commanded by wing motion patterns. C) Flexibility in the control of wing motion enables the bird to rapidly change the degree of feedforward and feedback control during a maneuvering sequence.

We found that the wing motion patterns among different demonstrations became more similar during evasive maneuvers (***𝒫𝓊-ℬ***) (wingbeats 2-4), compared with those during hover (***ℋ***). In both ***ℋ*** and ***𝒫𝓊-ℬ***, the body orientation also showed high similarity (or low variance). The variance of body orientation showed significant increases during the reorientation phase (***ℛ***) (wingbeats 5-8), while the wing motion similarity decreased (as the wing motion exhibited more diverse patterns). Towards the end of the reorientation phase (wingbeats 9-10), the body orientation becomes more similar, as the wing-motion similarity increased. Note that, the wing-motion similarity decreased again at the end of the reorientation (wingbeat 10) as the birds started to alter their wing motion to generate luminance-based responses to continue or terminate the escape.

## Discussion

### Rate-commanded maneuvers simplify flight control and require no active braking

We showed that the angular rates of hummingbird maneuvers are directly commanded by the changes in wing motion patterns (or WMPs), similar to the way the angular rate of an aircraft can be directly commanded by a human pilot (21, 22) via its electronic control system (32). For hummingbirds, such simplicity originates from its flight dynamics that map wing motion patterns to body angular motion on a wingbeat-by-wingbeat basis and can be considered as a form of physically-embodied intelligence (33). In other words, the flight dynamics effectively reduces to an approximate linear input-output kinematic relationship for a hummingbird’s flight control system (Flight dynamics, or a dynamical system in general can be approximated by using a kinematic relationship if it has a sufficiently low time constant or sufficiently fast response (34)). This trait likely alleviates the computational demand on flight control because it can reduce the complexity in an internal forward model. The internal forward model, which plays a crucial role in many aspects of motor control and learning, uses a copy of the motor command (i.e., an efference copy(35, 36)) to predict the outcome of motor control before sensory feedback is received, making control more efficient and accurate(36–40).

Rate-commanded maneuvers in hummingbirds also indicate that their wingbeat-averaged body angular rates are proportional to the changes in wing motions within the same wingbeat cycle. More specifically, within a single wing beat of an angular maneuver, the bird’s body experienced an acceleration followed by deceleration, resulting in a wingbeat-averaged “quasi-steady” body angular rate proportional to the magnitude of the WPM. This within-wingbeat oscillatory acceleration also led to a stepwise or impulsive change in angular rate, which was commonly observed in hummingbird maneuvers (13), including those studied here. Although the detailed mechanism relating such oscillatory acceleration to the rapid arrival to quasi-steady rate (within a wingbeat) remains to be elucidated (ongoing work of the authors), it is likely inherent to the oscillatory wing forces of comparable aerodynamic and inertial components (41), compounded with the low body inertia of hummingbirds. In addition, a stepwise or impulsive change in angular rate also suggests high within-wingbeat damping, either of passive nature, e.g., Flapping Counter Torque (FCT)(42, 43) inherent to flapping flight, or actively modulated via control. In addition, we observed that hummingbirds consistently flared their tail throughout the escape maneuvers, which likely also augmented the effective damping.

Another significant aspect of the rate-commanded maneuvers is that calliope hummingbirds did not have to actively brake from rapid body rotations. To decelerate from a rotation, they simply reduced the magnitude of the corresponding WMPs, and its body angular rate immediately followed (i.e., within the same wingbeat). In Figure 3A-i, during wingbeat 6-10, the bird’s body has roll acceleration (wingbeat 6-8) and deceleration (wingbeat 9-10). However, in Figure 2B-i, we observe that the asymmetry in wing kinematics responsible for body roll does not change its polarity (Note: the asymmetry between left-wing and right-wing kinematics creates a roll torque), only the magnitude of asymmetry changed i.e., the asymmetry first increased during wingbeat 6-8 and then decreased during wingbeat 9-10. This can be seen quantitatively in Figure 3A-ii, that the magnitude of WMP^***ℛ***^ did not reverse during the roll deceleration. Similarly, the magnitude of WMP^***𝒫𝓊-ℬ***^ did not reverse during pitch deceleration. This can be compared with a scenario when a human driver attempts to slow a car down by only releasing the gas pedal without hitting the brake; however, for the human driver, the car won’t be able to slow down fast enough (unless there is large damping, for example, if driving through deep water), so active braking is usually needed.

Despite the benefits to flight control, rate-commanded maneuvers likely demand a high energetic cost due to the putative high damping and high force generation, suggesting a tradeoff between the energetic cost and simplicity for control. Indeed, previous estimations show that the hummingbirds, when performing escape maneuvers, significantly increased both wingbeat frequency (which increases FCT damping (42)) and the pertinent mass-specific power (e.g., ∼4× higher power than those in hovering for Rivoli’s hummingbirds (14)).

### Hummingbirds excel in rapid flight control

This study quantified and reported how hummingbirds rapidly control a sequence of flight maneuvers, in terms of luminance-based responses and switching of maneuvering modules. Calliope hummingbirds were able to generate luminance-based responses in choosing ***𝒫𝒹-ℱ*** (category I) or ***𝒫𝓊-ℋ*** (category II) in less than 39±3 ms (bird 1) and 30±4ms (bird 2), and in choosing ***ℱ*** (category I) or ***𝒫𝓊-ℋ*** (category III) in less than 78±9ms (bird 1) and 91±20ms (bird 2). Hummingbirds also responded to the startling stimuli by initiating the escape and ***𝒫𝓊-ℬ*** in less than 60±17ms (bird 1) and 58±16 ms (bird 2). In addition, they were able to switch the maneuvering modules from ***ℋ*** → ***𝒫𝓊-ℬ*** → ***ℛ*** → ***𝒫𝒹-ℱ*** → ***ℱ*** →***𝒫𝓊-ℋ***, as many as 6 times in less than 250ms (category III, Figure 1D). To our best knowledge, no robotic flier has been described to date that is capable of generating a sequence of maneuvers of such celerity without losing stability.

We attribute the rapid flight control ability to the advanced visual and musculoskeletal systems of hummingbirds. Future comparative work would be useful to test whether aspects of these systems are shared, derived characteristics (synapomorphies) of hummingbirds, offering more effective, rapid control than in other bird species. Burst (anaerobic muscle capacity) is essential for the energetically demanding escape maneuver, but the use of creatine phosphate and glycolysis are shared features of vertebrate skeletal muscle. Moreover, hummingbird flight muscles exhibit homogeneity of fiber types similar to other small birds, and the myosin isoforms in their flight muscles are not unique (44, 45). Recent results (19) show that hummingbirds’ primary flight muscles have diverse effector capacities as they actuate all three-degree-of-freedom wing motion, and they experience a controlled tightening effect due to secondary muscles, which can also improve the ability of wing-motion control for hummingbirds. These traits can enable rapid force vectoring and transfer of muscle power of body maneuvers. Fast visual processing and vision-based responses are likely promoted by the hummingbirds’ capacity for gaze stabilization, which likely relies on the integrated global motion in the visual field and cervical reflexes. Hummingbirds have hypertrophied lentiformis mesencephali (LM) that encodes global optical flow information, and it responds more strongly to high-speed stimulus in near omnidirectionally fashion (46), which indicates that they have highly advanced visual sensory perception capability compared to other birds (47).

### Temporal variation in the cross-demonstration wing-motion similarity suggests varying degree of feedforward and feedback control during *ℋ* → *𝒫𝓊-ℬ* → *ℛ*

Vertebrates, including hummingbirds, produce rhythmic limb motions via motor programs that can be feedforward controlled, i.e., by a central controller generating a sequence of predetermined muscle activations, or be feedback controlled (35, 48), i.e., modulated by sensory feedback signals (e.g., visual or vestibular) measuring a bird’s flight states (49, 50). Hypothetically, feedforward control results in stereotyped wing motion patterns of high similarity among different escape flight demonstrations. In contrast, feedback control modulates the wing motions according to flight-state-dependent sensory feedback, which includes sources featuring uncertainty and noise, and it can add variability to the motor command generated by feedforward control alone, thereby decreasing the similarity of the resulting wing motion. We predicted the degree of feedforward vs feedback control of wing motion, by calculating the temporal variations in the cross-demonstration similarity of wing motion patterns throughout maneuvering modules ***ℋ*** → ***𝒫𝓊-ℬ*** → ***ℛ*** (Materials and Methods). Note that directly measuring the variability in muscle activation signals, which would provide precise information about motor command variability, is currently not feasible using electromyography (EMG) methods. This is due to the lack of knowledge in identifying the flight control muscles and the difficulty in EMG measurement during escape maneuvers in calliope hummingbirds.

Results showed that wing motions had higher similarity among demonstrations during evasive maneuvers (***𝒫𝓊-ℬ***) (wingbeats 2-4) than those in the rest of the escape, and therefore likely had a higher degree of feedforward control (Figure 4A-i). The evasive maneuver was the initial response to the startling stimulus and putatively generated large self-perturbation, which led to the observed high variability in the subsequent body motion (wingbeats 4-8, Figure 4A-ii). At the beginning of reorientation (***ℛ)***, the wing motion showed a decrease in similarity (wingbeats 5-8), suggesting that the birds modulated the wing motions as a result of increased degree of feedback control to correct the perturbations; therefore, we named this phase ***ℛ(ℱℬ)***. This prediction of increased feedback control is consistent with the reduced variability in body motion subsequently (Figure 4A-ii). However, towards the end of reorientation (wingbeats 9-10), wing motion showed increased similarity, indicating strengthened feedforward control (or weakened feedback control); we named this phase ***ℛ(ℱℱ)***.

In addition, hovering (***ℋ***, wingbeat 1) showed slightly lower similarity than those of evasive maneuvers, indicating that feedback was involved to stabilize hover, consistent with the previous finding that hovering flight is inherently unstable or weakly stable (51–53). Taken together, our similarity analysis for wing motion predicts that hummingbirds can flexibly alter the degree of feedback and feedforward control of WMPs 3 times in less than 160 ms (Figure 4 and Movie S7), i.e., ***ℋ(ℱℬ)*** → ***𝒫𝓊-ℬ (ℱℱ)***→ ***ℛ(ℱℬ)*** → ***ℛ(ℱℱ)***.

### Key traits of hummingbird agility

In summary, our results shows that hummingbird flight agility and stability are promoted by the following key traits. First, the ability to make vision-based decisions in the middle of rapid, highly-dynamic maneuvers, and the low latency in the visuomotor process (including both decision-making and neuromuscular wing-motion control). Together with proprioceptive feedback, they enable the rapid switch and mediation of wingbeat-by-wingbeat motion patterns during a maneuvering sequence. Second, the simplicity in the flight mechanics that maps wing motion patterns to body angular rates. This effectively reduces the “forward dynamics” for flight control (or the corresponding internal forward model (40, 54)) to an approximate input-output kinematic relationship (i.e., with a sufficiently low time constant (34)). Thus, hummingbird wing motion patterns directly rate-command body angular motion. Third, the flexibility in the control of wing motion enables changes in the degree of feedforward and feedback control during a maneuvering sequence. The combination of these traits, i.e., the *rapid, flexible control* of wing motion that *rate-commands* body angular motion, likely explains hummingbird-level agility and stability.

## Materials and Methods

### Experimental setup and protocols

We elicited escape flight in two calliope hummingbirds by startling each bird when it was hovering and feeding in a flight chamber (Figure 1A). We designed our experimental controls based on repeated trials on individual hummingbird subjects to identical startling stimuli and escape.

The initial condition during hovering and feeding was kept consistent throughout all the trials. After each escape flight was initiated, we switched off visible ambient LED light at various time instants, hence impeding the bird’s vision. This method enabled us to examine the hummingbirds’ responses to light removal, as well as the role of vision during the rapid escape flight, for example, testing whether vision was required in various phases of escape. The entire escape maneuver sequence was recorded using three highspeed monochrome cameras at 1000 fps under infrared light invisible to birds (57)(Figure 1A). we extracted body and wing kinematics using digitization (58) of anatomical landmarks (*SI Appendix*, Figure S1). More details on the bird’s preparation, experimental setup, data post-processing, and kinematic extraction are in the *SI Appendix*.

### Statistical analysis

#### Calculation of critical light removal time instant 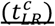

We defined 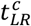 as the critical light removal time instance. The visual input to the birds at 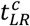was hypothesized to influence the birds’ responses, i.e., whether the bird’s vision was blocked or not separated categories II and III (Figure 1). Logistic regression was performed on the *t*_*LR*_ and escape categories to estimate 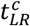(see *SI Appendix*, Figure S2 for details).

#### Relationship between t _SH_ and t_LR_

We performed hypothesis testing to establish the relationship between *t*_*SH*_ and *t*_*LR*_ and showed that *t*_*SH*_ was independent of the *t*_*LR*_ in category II, and *t*_*SH*_ varied linearly with *t*_*LR*_ in category III (Figure 1D, details in *SI Appendix*).

### Analysis of Wing Motion Primitives (WMPs)

In robotics, Motion Primitives (MPs) are commonly used in modular movement generators (59–61), and in modeling human motion for robot learning from demonstrations (62, 63). In this work, we modeled hummingbird wing motion patterns during escape maneuvers as Wing Motion Primitives (WMPs), similar to kinematic motion primitives (kMPs)(20).

#### Extraction of WMPs and calculation of WMP magnitude for individual wing beats

To extract the maneuvering WMPs during the ***ℋ*** → ***𝒫𝓊-ℬ*** → ***ℛ*** that are common to all demonstrations, we first assumed that all the individual demonstrations emerged as noisy observations of a single “hidden” nominal trajectory, and we estimated this trajectory as the intended trajectory among all demonstrations (23, 64, 65), following the methods used in *Abbeel,P*., *et al* (64, 65) (for details on the estimation of intended trajectory see *SI Appendix*). The hover Wing Motion Primitive (WMP^***ℋ***^) was defined as the average of all the hovering wingbeat motion patterns (N=40 wingbeats for bird 1 and N=36 for bird 2). It is then assumed that the maneuvering WMPs would be very different from the hovering WMPs. We then measured the statistical difference using Hellinger distance (24) between the WMP^***ℋ***^ and each maneuvering wingbeat from the estimated intended trajectory,

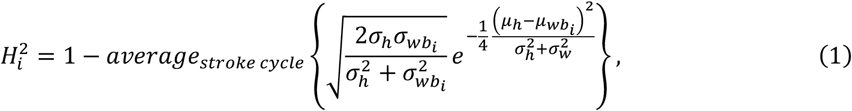

where 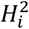 is the squared Hellinger distance between two normal distributions hover wingbeat ∼ 𝒩 (*μ*_*h*_, *σ*_*h*_) and *i*^th^ maneuvering wingbeat 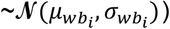. The maneuvering WMPs were then defined at the local maxima and we named WMPs according to the maneuvering modules that they correspond to (WMP^***𝒫𝓊-ℬ***^: the WMP for evasion, occurred at the 3^rd^ wingbeat, and WMP^***ℛ***^: WMP for reorientation, occurred at the 8^th^ wingbeat, Figure 2B).

We assumed that individual wingbeats were composed of WMP^***ℋ***^, WMP^***𝒫𝓊-ℬ***^, and WMP^***ℛ***^, and to find the wingbeat-by-wingbeat magnitude of each WMP during the maneuvers, we reconstructed individual wingbeats based on a weighted linear combination of WMPs (20),

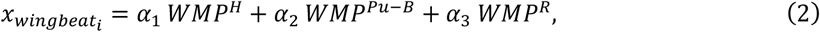

where *α*_*j*_ is the weight of each WMP, which represents the WMP magnitude in *i*^*th*^ wing beat 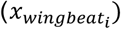 (Figure 3A-ii). We then calculated the cross-correlation (*xcorr* function in MATLAB, MathWorks, Inc., Natick, MA) of WMPs magnitude with respective cycle-averaged angular velocity and angular acceleration (Figure 3B) (see *SI Appendix* for details).

#### Similarity analysis of wing kinematics during escape maneuver

We assumed that feedforward flight control results in wing motion patterns of high similarity (or less variability) among multiple escape flight demonstrations, while feedback control results in low similarity (or more variability) due to modulation of wing motion driven by sensory feedback that is flight state-dependent (details on the assumption is explained in *SI Appendix*). We measured the wing kinematic similarity among different escape flight demonstrations using the Coefficient of Multiple Correlations (CMC) (Equation 3). CMC is widely used in gait cycle analysis to measure the similarity amongst multiple demonstrations (30, 31, 66–70).

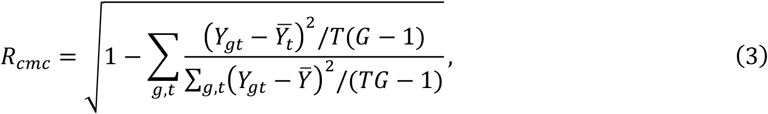

where *T* and *G* are the number of time instants in each demonstration and the number of demonstrations, respectively, *Y*_*gt*_ is the wing Euler angle in consideration at time *t* of *g*^*th*^ test demonstrations, 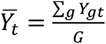 is the mean across all test demonstrations *a*, and 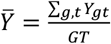 is the grand mean across all test demonstrations *a* and timepoints *t* (See *SI Appendix* for details).

We calculated *R*_*cmc*_ for wing deviation angles for a two-wingbeat sliding window (∼36 ms), which slides from the 1^st^ wingbeat of all escape demonstrations to subsequent wingbeats with an increment of half wingbeat (See *SI Appendix* for details). We chose only the wing deviation angle for similarity calculation because hummingbirds manipulate primarily the wing deviation angle for control (71), while stroke angle is set by flapping nature, and wing pitching is primarily passive in hummingbirds (41).

### Nomenclature

|  |  |
| --- | --- |
| $\mathcal{H}$ | Hover module |
| $\mathcal{Pu-B}$ | Evasive module (*Pitch up* to *Backward* flight) |
| $\mathcal{R}$ | Reorientation module (*Roll* to reorient) |
| $\mathcal{Pu-H}$ | *Pitch up* to *Hover* module |
| $\mathcal{Pd-F}$ | *Pitch down* to *Forward* flight module |
| $\mathcal{F}$ | Forward flight Module |
| $t_{LR}$ | Light removal time (the time from initiation of escape flight to the moment when the visible light was removed) |
| $t_{LR}^c$ | Critical light removal time |
| $t_{SH}$ | Time instant to stop and hover, which marks the end of the escape flight (the time from initiation of escape flight to the end of pitch up to hover ($\mathcal{Pu-H}$) module) |
| $WMP^{\mathcal{H}}$ | Wing motion primitive for hover |
| $WMP^{\mathcal{Pu-B}}$ | Wing motion primitive for evasion (*Pitch up*) |
| $WMP^{\mathcal{R}}$ | Wing motion primitive for reorientation (*Roll*) |
| $R_{cmc}$ | Coefficient of Multiple Correlations (CMC) |

## Data Availability

All data related to this work are available on Dryad. (https://datadryad.org/stash/share/YagXzSdL06FbmbDPDYXjKDpR8uQvimgaHAxvJ6EqrK8).

## Footnotes

1 In the experiment there was a hard separation of *t*_*LR*_ between category II and III, and from the logistic regression the 68% bound window is less than 1 ms (least count of measurement). Hence 68% bound is not reported here.

