## Supplementary material for "Hummingbirds rapidly respond to the removal of vision and control a sequence of rate-commanded maneuvers in milliseconds": SI Appendix

**This PDF file includes:**

Supporting Information Text
Figures S1 to S12
Tables S1 to S2
Legends for Movies S1 to S7
SI References

**Other supporting materials for this manuscript include the following:**

Movies S1 to S7

### Supporting Information Text

#### Experiments on Hummingbird Escape Maneuvers

We studied escape flight in two male calliope hummingbirds (Selasphorus calliope) (body mass of 2.1 g and 2.4 g). We removed the bird's vision at various instants during the escape maneuvers and studied the roles of visual feedback, decision-making, and flight control.

Bird preparation and protocols

Birds were collected and temporarily housed during May and June 2020 in Missoula, MT (Missoula County), USA. In the laboratory, birds were housed individually in 1×1×1$m$ cages and fed a 50:50 mixture of 20% sucrose solution and Nectar Plus© *ad libitum* using a Dr. JB's hummingbird feeder. Both study subjects maintained mass during captivity, confirming that the feeder solution adequately supplied essential dietary needs. The birds were released at the point of capture after experiments were complete. Scientific collecting was under permits from the US Fish and Wildlife Service and Montana Department of Fish, Wildlife, and Parks. All protocols were approved by the University of Montana Institutional Animal Care and Use Committee (IACUC).

Experimental setup and protocols

We performed the experiments in a long corridor of size 5.6m $\times$1.8m $\times$1.8m, which was sufficiently large to provide an escape path to hummingbirds comparable to naturalistic settings. For the experiment, a single bird was introduced into the corridor, and an experimenter was present behind a feeder syringe inside the corridor. Then, the experimenter waited behind the feeder for the bird to hover and feed on the sucrose solution in the feeder, and applied a startling stimulus using the rapid movement of a red board (23 cm $\times$ 32 cm) towards the bird. A proximity sensor was used to detect the movement of the red board, based on which it started the high-speed camera recording, and switched off the power to visible LED lights (thereby removing hummingbird vision) at an array of specified time delays using a microcontroller (Arduino-Uno, Arduino, Somerville, MA, USA) and a nominally-closed switch (NC switch) (Figure 1A).

High-speed video recording and lighting conditions

The birds' behaviors were recorded using three shutter-synchronized high-speed monochrome cameras (one FASTCAM NOVA S6 model with 28 mm zoom, and two FASTCAM Mini UX 100 model with 18 mm zoom and 20 mm zoom, Photron Inc., San Diego, CA, USA) at a 1000 fps. Six IR (Infrared) lamps (wavelength of 850 nm), which were invisible to hummingbirds(1), were used for camera recording. Ambient light visible to hummingbirds was provided by LED lights. All the other possible light sources in the experiment room were covered with opaque black sheets. The luminous flux in the corridor was measured using a digital light meter (ET130, Klein Tools, Inc., Lincolnshire, IL, USA); during the experiment, it was ~110 fc with the LED lights on, and ~2 fc with LED lights off in front of the IR lamps (at 1 cm distance from the lamps), and 0.00 fc elsewhere including the escape flight region.

#### Data Processing

Data for post-processing

During the experiment, a total of 50 experimental trials were recorded for each bird. Both birds demonstrated both left and right turn (with combined roll and yaw) at the initiation of the escape flight (bird 1: 36/50 (left) and 14/50 (right); bird 2: 31/50 (left) and 19/50 (right)). The observed escape behaviors of birds between left and right turns were mirrored versions of each other. For post-processing and flight kinematics analyses, we only used the trials with left turns and excluded those in which the bird was spooked by involuntary subtle movements of the experimenter instead of startling stimuli or started the escape flight at a location different than the feeder (because the bird got spooked and at the time of startling stimuli bird was not hovering at the feeder). As a result, a total of 22 trials for bird 1 and 21 trials for bird 2 were post-processed.

In the calculation of time to stop and hover ($t_{SH}$), escape flights with both left and right turns were used (45 trials for bird 1, and 32 trials for bird 2, we again excluded those trials in which birds did not execute a complete escape flight or started the escape flight at a location different than the feeder).

Post-processing and filtering of data

We digitized ten points on a bird (two on the beak, four on the body, and four on the wings, Figure S1) at 1000 fps using direct linear transformation calibrations (DLTdV8)(2), which was obtained using easyWand5(2) with wand and ball drop. The digitized point coordinates were filtered at 40 Hz for body points and 400 Hz for wing points (more than six times their wingbeat frequency) using a sixth-order digital low-pass Butterworth filter to remove digitization noises. The data were then interpolated at 0.1 ms intervals using a cubic spline.

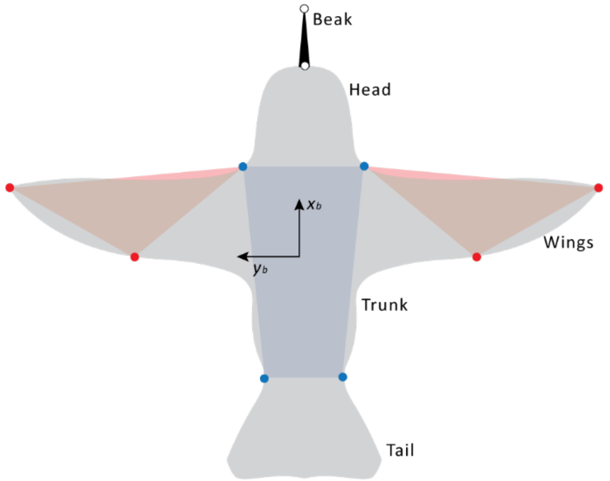


**Figure. S1. Anatomical marks on the bird for extracting body and wing kinematics.** 10 points on the bird (4 on the body (blue), 4 on the wings(red), and 2 on the beak(white)) were digitized and used for calculating the body and wing kinematics.

Coordinate systems and definitions for body and wing kinematics

To describe a hummingbird's body and wing kinematics, a global coordinate frame $\{X_g, Y_g, Z_g\}$ and a body coordinate frame $\{x_b, y_b, z_b\}$ were defined. The global positive Z-axis was aligned with negative gravity (defined by the direction of the ball drop), the X-axis was defined as the initial hummingbird heading before the start of the escape maneuver, and the Y-axis was defined according to the right-hand rule. The global origin was placed at the feeder tip. The body frame $\{x_b, y_b, z_b\}$ was attached to the bird's body at a point defined by the averaged shoulder and hip points (blue points, Figure S1), $x_b$ defined as the body's long axis (tail to head), $y_b$ as the body's transverse axis (right to left), and $z_b$ as the body's normal axis (ventral to dorsal). Body orientation w.r.t. the global frame was defined by rotating the global frame to the body frame according to Euler angles of the Z-Y-X (yaw-pitch-roll) convention. Body angular rates were defined with respect to the body's fixed frame $\mathcal{F}_b = \{x_b, y_b, z_b\}$ (Figure 2A), with body roll as rotation about the $+x_b$ axis, pitch about the $-y_b$ axis, and yaw about the $+z_b$ axis. Wing kinematics relative to the body are defined by Euler angles of Z-X-Y convention (Stroke-Deviation-Pitch) (see definitions in Figure 2A).

Definition of maneuvering modules

In Figure 1B, the maneuvering modules from hover to reorientation (i.e. $\mathcal{H} \rightarrow \mathcal{P}u\text{-}\mathcal{B} \rightarrow \mathcal{R}$) were qualitatively defined based on observed body rotation. While the segmentation points between $\mathcal{R} \rightarrow \mathcal{P}d\text{-}\mathcal{F}$ or $\mathcal{R} \rightarrow \mathcal{P}u\text{-}\mathcal{H}$ were defined by the binary change in wing motion between $\mathcal{P}d\text{-}\mathcal{F}$ and $\mathcal{P}u\text{-}\mathcal{H}$. which occurs at the beginning of the downstroke of the 10th wingbeat (152±3 ms for bird 1 and 147±2 ms for bird 2).

### Statistical Analysis

Calculation of critical light removal time instant ($t_{LR}^{c}$) for Category II and Category III

The $t_{LR}^{c}$ was calculated as the critical light removal time instant that separated categories II and III (Figure 1D). In category II (Figure 1B-ii), where the visible light was removed before $t_{LR}^{c}$ during the escape, the bird did not initiate the ($\mathcal{Pd}$-$\mathcal{F}$) after ($\mathcal{R}$) instead it pitched up and returned to hover ($\mathcal{Pu}$-$\mathcal{H}$). While in category III (Figure 1B-iii), where the visible light was removed after $t_{LR}^{c}$ during the escape, the bird did initiate ($\mathcal{Pd}$-$\mathcal{F}$) but forward flight ($\mathcal{F}$) was terminated after a delay following the instants of light removal, and then the bird performed $\mathcal{Pu}$-$\mathcal{H}$ to hover in the dark.

To calculate $t_{LR}^{c}$, we performed logistic regression (glmfit function in MATLAB, MathWorks, Inc., Natick, MA) for light removal timing $t_{LR}$ and escape categories (with binary values 0 for category II and 1 for category III) (Figure S2). Logistic regression provided the probability of a demonstration being in category II or III. The probability of a demonstration belonging to Category III is given by $\pi$ (Equation S1) and to Category II is given by $(1-\pi)$. We defined the time instant at which probability crosses 0.5 level as the "critical light removal time" $t_{LR}^{c}$ (Bird 1: 113 ms, and Bird 2: 116 ms).

$$\pi = \frac{exp(\beta_0 + \beta_1 t_{LR})}{1 + exp(\beta_0 + \beta_1 t_{LR})}, \qquad (S1)$$

where $\pi$ is the probability of a demonstration belonging to category III as a function of light removal time $t_{LR}$, $\beta_0$ and $\beta_1$ are the optimized parameters for logistic regression (Table S1). $-\beta_1/\beta_0$ is the time instant where the probability crosses 0.5 (i.e., $t_{LR}^{c} = -\beta_1/\beta_0$) and $\beta_1$ determines the softness of the classification boundary (a high value means hard separation while a low value means soft separation). For Bird 1, the data formed a hard boundary meaning all the demonstrations for which $t_{LR} < t_{LR}^{c} = 113\,ms$, belonging to category II while for $t_{LR} > t_{LR}^{c} = 113\,ms$ they belong to category III. On the other hand, bird 2 data formed a soft boundary with some overlapping data (Figure S2). A 0.68 probability window (1 s.d probability) around the critical light removal time $t_{LR}^{c}$ was found to be 116 ± 3 ms.

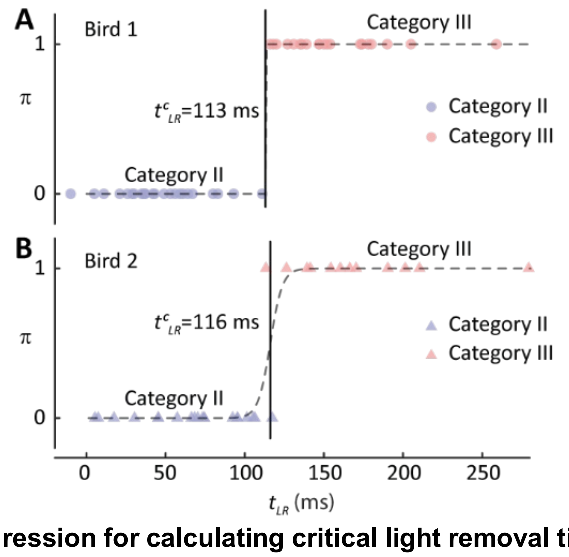


**Figure. S2. Logistic regression for calculating critical light removal time instant.** The dashed curves represent the probability of an escape demonstration to be in category III ($\pi$) or category II ($1-\pi$) versus light removal time $t_{LR}$. The time at which the probability crosses 0.5 value is the critical light removal time $t_{LR}^{c}$.

**Table S1. Optimized parameters from logistic regression.**

| | $\beta_0$ | $\beta_1$ | $t_{LR}^{c} = -\beta_1/\beta_0$ |
|---|---|---|---|
| Bird 1 | -1449.10 | 12.76 | 113.56 |

| | | | |
|---|---|---|---|
| Bird 2 | -30.36 | 0.26 | 115.75 |

<u>Hypotheses testing for the relationship between $t_{SH}$ and $t_{LR}$</u>

In all demonstrations, after the visible light was removed at $t_{LR}$, the bird eventually performed $\mathcal{Pu}$-$\mathcal{H}$ and hover at a time instant to stop and hover ($t_{SH}$). The relationship between $t_{SH}$ and $t_{LR}$ are shown in Figure 1D, based on which we hypothesized that $t_{SH}$ was independent of the $t_{LR}$ in category II, and $t_{SH}$ varied linearly with $t_{LR}$ in category III. To test these hypotheses, the following statistical tests were performed.

For category II:

***H0***: There is no statistically significant relationship between $t_{SH}$ and $t_{LR}$ in demonstrations of category II, i.e., $t_{SH}$ was independent of the $t_{LR}$ in category II.

***H1***: $t_{SH}$ is dependent on $t_{LR}$ in demonstrations of category II.

The P-value for the linear term in the linear fit model between $t_{SH}$ and $t_{LR}$ for bird 1 and bird 2 were found to be 0.28141 and 0.36368, respectively. Hence at a significance level of alpha=5%, we cannot reject the null hypothesis. For category II, $t_{SH}$ = 193 ± 5 ms for bird 1 and $t_{SH}$ = 199 ± 10 ms for bird 2.

For category III:

***H0***: There is no statistically significant relationship between $t_{SH}$ and $t_{LR}$ in demonstrations of category III, i.e., $t_{SH}$ was independent of the $t_{LR}$ in category III.

***H1***: $t_{SH}$ is dependent on $t_{LR}$ in demonstrations of category III.

The P-value for the linear term in the linear fit model between $t_{SH}$ and $t_{LR}$ for bird 1 and bird 2 were found to be $7.06\times10^{-6}$ and $4.26\times10^{-15}$, respectively. Hence at a significance level of alpha=5%, we reject the null hypothesis and conclude that there is a linear relation between $t_{SH}$ and $t_{LR}$, with a slope of 1.05 ± 0.05 and 1.04 ± 0.12 for bird 1 and bird 2, respectively.

**Intended Trajectory during Escape Maneuvers**

This section describes the method to obtain the intended trajectory, which was used to extract Wing Motion Primitives (WMPs, described in the following section). We first assumed that the bird was the expert(3) executing a single intended trajectory during different demonstrations. Individual demonstrations appeared warped in time, i.e., they exhibited similar patterns of body and wing motions, but appear dephased in time(4). Moreover, these demonstrations exhibited uncertainties (which likely arose from both biological and physical processes) added to the intended trajectory, in other words, the individual escape demonstrations can be considered as noisy observations of the hidden, intended trajectory. Therefore, we used the time-aligned (dynamic time-wrapped) average of all the demonstrations as the estimate of the underlying intended escape trajectory (Figure 2B-i).

Specifically, for the time-aligned estimate, we assumed that the measured escape trajectories were noisy observations $\boldsymbol{Y}^k$ (with $N^k$ number of observations (time instances), k is the index for each demonstration, a total of $M$ demonstrations) from the identical intended trajectory, modeled by hidden states $\boldsymbol{Z}$ (with $N_s$ number of states), with additive noise ($\boldsymbol{\omega}^y$). A generative model (Equation S2) represents each demonstration $y_j^k$ as a set of independent “observations” of the hidden intended trajectory $z_{\tau_j^k}$. $\tau_j^k$ is the time index in the hidden trajectory to which the observation $y_j^k$ is mapped(3, 5) (Figure S3).

$$y_j^k = z_{\tau_j^k} + \omega_j^{(y)} \quad \omega_j^{(y)} \sim N\left(0, \Sigma^{(y)}\right), \qquad j = 1,2 \dots . N^k \qquad (S2)$$

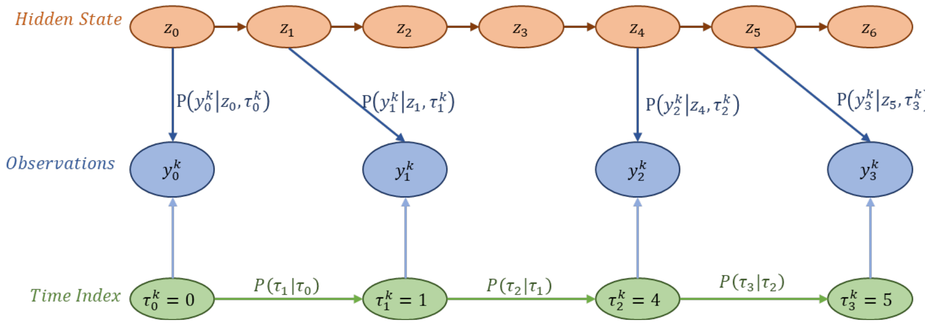


**Figure. S3. Graphical model representing intended trajectory assumptions**. Blue nodes are observed (kinematics data of individual demonstrations $\boldsymbol{Y}^k$), orange nodes are hidden states (intended trajectory $\boldsymbol{Z}$), and green nodes are time indexes ($\boldsymbol{\tau}^k$) that map observation onto the intended trajectory**.**

To accommodate the time wrapping, it was assumed that the observed trajectories were subsampled versions of the hidden trajectory i.e. $N_s > N^k$. Time shifts during time alignment according to the following model (Equation S3) with a maximum allowed shift to be $\delta\tau_{max}$

$$P\left(\tau_{j+1}^k \middle| \tau_j^k\right) = \begin{array}{ll} d_{\delta\tau}^k & if\ \tau_{j+1}^k - \tau_j^k = \delta\tau \ \text{ for } \delta\tau = 1,2, \ldots \delta\tau_{max} \\ 0 & if\ \tau_{j+1}^k - \tau_j^k = \delta\tau \ \text{ for } \delta\tau > \delta\tau_{max} \end{array}, \qquad (S2)$$

for our work we chose $\delta\tau_{max} = 5$ ms and $d_{\delta\tau}^k = \frac{1}{\delta\tau_{max}}$ $\forall\ \delta\tau = 1,2, \ldots \delta\tau_{max}$.

The time alignment and state estimation were done iteratively where an estimate of the trajectory $(z_{\tau_j^k})$ was found for a given time alignment index $(\tau_j^k)$ and for that estimated trajectory a new alignment index was calculated to maximize the likelihood of the demonstrated trajectories.

<u>Time alignment Algorithm</u>

Assume initial $\boldsymbol{\tau}^k$ for all $k$ demonstrations. Estimate hidden states $\boldsymbol{Z}$ that maximize the likelihood of observing $\boldsymbol{Y}^k$ observations given $\boldsymbol{\tau}^k$ (MLE estimate). In the next step, find $\boldsymbol{\tau}^k$ that maximizes the likelihood of observing $\boldsymbol{Y}^k$ observations for a given $\boldsymbol{Z}$ (Equation S4)

$$\boldsymbol{\tau}^k = \underset{\boldsymbol{\tau}^k}{arg\,max} \sum_{k=0}^{M-1} \sum_{j=0}^{N^k-1} \left\{ L\left(y_j^k \middle| z_{\tau_j^k}, \tau_j^k\right) + L\left(\tau_j^k \middle| \tau_{j-1}^k\right) \right\}, \qquad (S4)$$

where, $L\left(y_j^k \middle| z_{\tau_j^k}, \tau_j^k\right)$ is the likelihood of observing $y_j^k$ given $z_{\tau_j^k}$ and $\tau_j^k$; $L\left(\tau_j^k \middle| \tau_{j-1}^k\right)$ is the likelihood for the time shift.

---

**Pseudo Code:** Finding the time index that maximizes the likelihood(3, 6)

---

**Input:** Observation data $\boldsymbol{Y}^k$ $(N^k\ observations)$; Number of states $N_s$ for intended trajectory
**Initialize**: Assume initial time index $\boldsymbol{\tau}^k$
**Repeat:**

Calculate the Covariance matrix $\boldsymbol{R}$ and estimate $\boldsymbol{Z}$ ($N_s$ number of states) from the observation data $\boldsymbol{Y}^k$ and time index $\boldsymbol{\tau}^k$

Formulate $\boldsymbol{Q}_p^k$ matrix for each demonstration, with

$$\boldsymbol{Q}_p^k(1,j) = L\left(y_1^k \middle| z_{\tau_1^k}, \tau_1^k = t_1, \boldsymbol{R}\right) + L(\tau_1^k = t_1)$$

$$\boldsymbol{Q}_p^k(i,j) = L\left(y_i^k \middle| z_{\tau_i^k}, \tau_i^k = t_i, \boldsymbol{R}\right) + \max_{t'}\{L(\tau_i^k = t_i \,|\tau_{i-1}^k = t_{i-1}) + \boldsymbol{Q}_p^k(i-1, t_{i-1})\}$$

$\boldsymbol{Q}_t^k$ matrix is formed as

$$\boldsymbol{Q}_t^k(i,j) = index\ value\ \left(\max\left(L(t_j|t_i) + \boldsymbol{Q}_p^k(i,t_i)\right)\right)$$

The updated time index $\boldsymbol{\tau}^k$ for the $k^{th}$ demonstration is given by

$$\tau_{N^k}^k = \boldsymbol{Q}_t^k(N^k - 1, N_s)$$

$$\tau_i^k = \boldsymbol{Q}_t^k(i, \tau_{i+1}^k)$$

**End**: Convergence

---

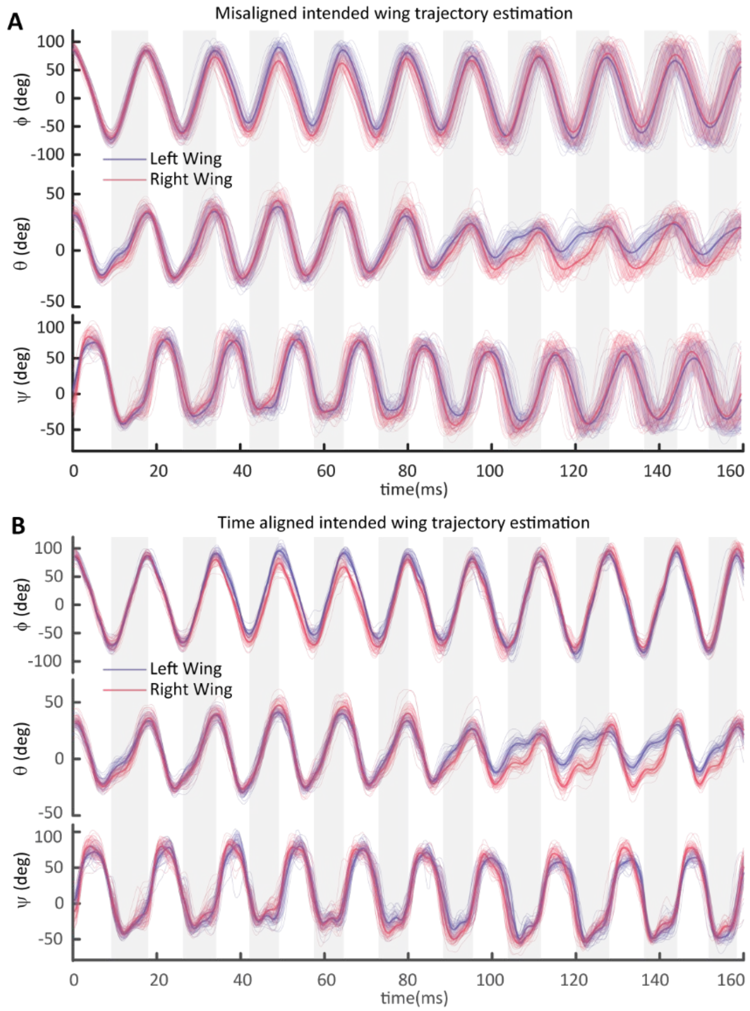


**Figure. S4. Comparison of unaligned and time-aligned estimation of intended trajectory.** Estimate of Intended trajectory (thick line) during $\mathscr{H} \rightarrow \mathscr{Pu}\text{-}\mathscr{B} \rightarrow \mathscr{R}$ maneuvering modules (A) from unaligned and (B) time-aligned kinematics from individual demonstrations (thin lines), the shaded areas enclosing the curves indicate ±1 s.d. (22 demonstrations).

The time-alignment method improved the accuracy of the estimation of the intended trajectory. The time-aligned estimates have tighter standard deviations as compared to the unaligned estimate (Figure. S4). The mean standard deviation (S.D.) in wing angle estimation using both methods is given in Table S2.

**Table S2. Averaged standard deviation in wing angle estimation**

| | Unaligned estimate (Average s.d.) | Time-aligned estimate (Average s.d.) |
|---|---|---|
| Stroke angle (ϕ) | 10 deg (bird 1), 28 deg (bird 2) | 09 deg (bird 1), 32 deg (bird 2) |
| Deviation angle (θ) | 07 deg (bird 1), 13 deg (bird 2) | 06 deg (bird 1), 11 deg (bird 2) |
| Pitch angle (ψ) | 13 deg (bird 1), 26 deg (bird 2) | 11 deg (bird 1), 25 deg (bird 2) |

**Analysis of Wing Motion Primitives (WMPs)**

In robotics, Motion Primitives (MPs) are commonly used in modular movement generators(7–9), and in modeling human motion for robot learning from demonstrations(10, 11). In this work, we described the wing trajectory during the escape maneuver using Wing Motion Primitives (WMPs) similar to kinematic motion primitives (kMPs)(12).

Extraction of WMPs

For extracting the WMPs from the wing kinematics of all the demonstrations, we first found the intended trajectories during $\mathcal{H} \rightarrow \mathcal{Pu}\text{-}\mathcal{B} \rightarrow \mathcal{R}$ modules of the escape maneuver (see above section). We then defined hover Wing Motion Primitive (WMP$^{\mathcal{H}}$) based on averaging the wingbeat motion patterns preceding the evasion (N=40 wingbeats for bird 1 and N=36 for bird 2). To obtain the maneuvering WMPs during the $\mathcal{H} \rightarrow \mathcal{Pu}\text{-}\mathcal{B} \rightarrow \mathcal{R}$, we measured the statistical difference (Hellinger distance(13)) (Equation S5) between the WMP$^{\mathcal{H}}$ and each maneuvering wingbeat from the estimated intended trajectory. Each wingbeat is represented by a concocted vector of wing angles $[\phi_{LW}; \theta_{LW}; \psi_{LW}; \phi_{RW}; \theta_{RW}; \psi_{RW}]$.

$$H_i^2 = 1 - average\left\{\sqrt{\frac{2\sigma_h\sigma_{wb_i}}{\sigma_h^2 + \sigma_{wb_i}^2}}\, e^{-\frac{1}{4}\frac{\left(\mu_h - \mu_{wb_i}\right)^2}{\sigma_h^2 + \sigma_w^2}}\right\}, \qquad (S5)$$

where, $H_i^2$ is the squared Hellinger distance between two normal distributions hover wingbeat $\sim\mathcal{N}(\mu_h, \sigma_h)$ and $i^{th}$ maneuvering wingbeat $\sim\mathcal{N}(\mu_{wb_i}, \sigma_{wb_i})$), $\mu_{hover}$ and $\sigma_h$ are the mean and standard deviation of the hovering wingbeat estimate, respectively; $\mu_{wb_i}$ and $\sigma_{wb_i}$ are the mean and standard deviation of the $i^{th}$ maneuvering wingbeat from the estimated intended trajectory. The term inside the bracket is for each time instance during the wingbeat which is then averaged over the stroke cycle. When the $i^{th}$ maneuvering wingbeat is the same as the hovering wingbeat then $distance_i = 0$ and in case $i^{th}$ wing beat is different from hovering then $distance_i \sim 1$.

The maneuvering WMPs were defined at the local maxima (WMPs for evasion (WMP$^{\mathcal{Pu}\text{-}\mathcal{B}}$) at the 3rd wingbeat and WMP for reorientation (WMP$^{\mathcal{R}}$) at the 8th wingbeat, Figure 2B). These wingbeats also correspond to the maximum pitching and maximum rolling rate, respectively.

Wing trajectory reconstruction from WMPs

To estimate the magnitude of each WMP in individual wing beat patterns of escape demonstrations, every wingbeat was reconstructed(12) using three derived WMPs (WMP$^{\mathcal{H}}$, WMP$^{\mathcal{Pu}\text{-}\mathcal{B}}$, and WMP$^{\mathcal{R}}$). Reconstruction is based on a weighted linear combination of WMPs,

$$x_{wingbeat_i} = \alpha_1\, WMP^H + \alpha_2\, WMP^{Pu-B} + \alpha_3\, WMP^R, \qquad (S6)$$

where WMP$^{\mathcal{H}}$, WMP$^{\mathcal{Pu}\text{-}\mathcal{B}}$, and WMP$^{\mathcal{R}}$ represent WMPs for $\mathcal{H}$, $\mathcal{Pu}\text{-}\mathcal{B}$, and $\mathcal{R}$, respectively (each WMP is a concocted vector of wing angles $[\phi_{LW}; \theta_{LW}; \psi_{LW}; \phi_{RW}; \theta_{RW}; \psi_{RW}]$), and $\alpha_j$ is the weight of each WMP, which represents the WMP magnitude in $i^{th}$ wing beat ($x_{wingbeat_i}$)(Figure 3A-ii). To calculate $\alpha_j$, we solved the minimum least-squares problem.

Relation of WMP magnitude with angular rate

We fitted a linear model for angular rates with WMP magnitude (Equation S7). We found that WMP$^{\mathcal{Pu}\text{-}\mathcal{B}}$ magnitude shows a proportional trend with the pitch rate and an inverse trend with the roll rate (Figure 3C-i). Similarly, the WMP$^{\mathcal{R}}$ magnitude shows a proportional trend with the roll rate and an inverse trend with the pitch rate (Figure 3C-ii).

$$w_{pitch,roll} = \left(a + b\,\alpha_{Pu-B,R}\right)\left|w_{pitch,roll}\right|_\infty, \qquad (S7)$$

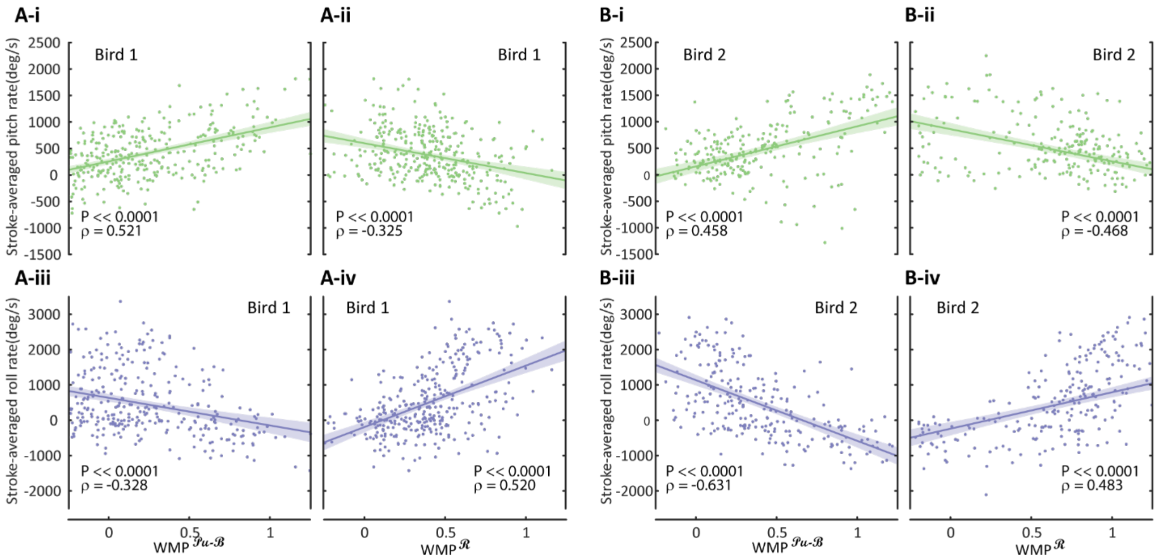


**Figure. S5. The relationship between stroked-averaged angular rates and WMPs magnitude.** Relationship graphs of (A) bird 1 and (B) bird 2. (A-i, B-i) Pitch rate vs WMP$^{\mathscr{Pu}\text{-}\mathscr{B}}$ magnitude, (A-ii, B-ii) Pitch rate vs WMP$^{\mathscr{R}}$ magnitude, (A-iii, B-iii) Roll rate vs WMP$^{\mathscr{Pu}\text{-}\mathscr{B}}$ magnitude, and (A-iv, B-iv) Roll rate vs WMP$^{\mathscr{R}}$ magnitude. The solid lines represent linear fit to data and the shaded area represents the confidence interval at a significance level of 0.05. Pearson's linear correlation coefficient (ρ) and P-value between each angular rate and WMP are shown at the bottom of each graph.

<u>Details on wing similarity calculation</u>

To calculate the similarity of wing motion patterns, we used the following procedure. First, each wingbeat within an escape maneuver demonstration $\mathscr{H} \rightarrow \mathscr{Pu}\text{-}\mathscr{B} \rightarrow \mathscr{R}$ was numbered, with the last wingbeat at hover ($\mathscr{H}$) preceding the escape manuever numbered as 1. Then, we implemented a two-wingbeat sliding window, approximately 36 ms in duration, which moved from wingbeat #1 to subsequent wingbeats with a half-wingbeat increment (Figure S6). This sliding window allowed us to assess the similarity of wing motion among all the demonstrations within the defined window.

The selection of the window size was based on empirical considerations. We aimed to keep the window narrow enough to capture within-wingbeat differences while ensuring it was large enough to detect any local frequency changes. It is worth noting that a single wingbeat window would not have captured the local phase shift due to frequency variations effectively. Hence, we opted for a 2-wingbeat window, which enabled us to also capture the dissimilarities caused by temporal variations.

While the standard deviation does provide a measure of variability by quantifying the spread or dispersion at a specific point in the wing motion, it alone does not offer a comprehensive understanding of the overall similarity or dissimilarity of the wing motion patterns. Therefore, we incorporated the sliding window approach with the Coefficient of Multiple Correlation (CMC) (14–20) analysis to obtain a more comprehensive assessment of the similarity among the wing motion patterns.

By implementing this specific procedure, we aimed to capture the temporal variations, local frequency changes, and overall similarity or dissimilarity of the wing motion patterns during the escape maneuvers, providing a more detailed understanding of the dynamics involved.

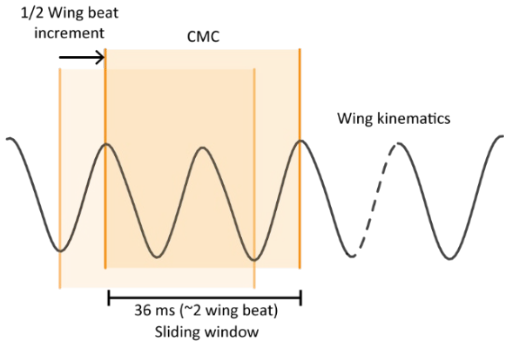


**Figure S6. Similarity calculation window.** Two wingbeats sliding window with an increment of a half wing beat for calculation of similarity of wing kinematics amongst different demonstrations of escape maneuver.

### Supplementary Texts

#### No observed learning and adaptation during the experiment

For the experiment, we ensured that the initial conditions during hovering and feeding were consistent across all trials. The hummingbird's escape maneuver is a natural "fight-or-flight" response to avoid danger that is likely executed with maximal performance, therefore reducing the possibility of adaptation or learning. To further minimize the effects of adaptation and learning, we designed the flight chamber to be sufficiently large to allow for natural response. We also allowed hummingbirds to regularly feed without startling stimuli and we applied waiting time between the trials, during which the hummingbirds were able to calmly perch. As a result, we did not observe major adaptations in the birds' behaviors as a function of time or trial number during the study, and their nominal responses without light removal were stereotypical (Figure S7). This result is consistent with previous studies on hummingbird escape maneuvers(21).

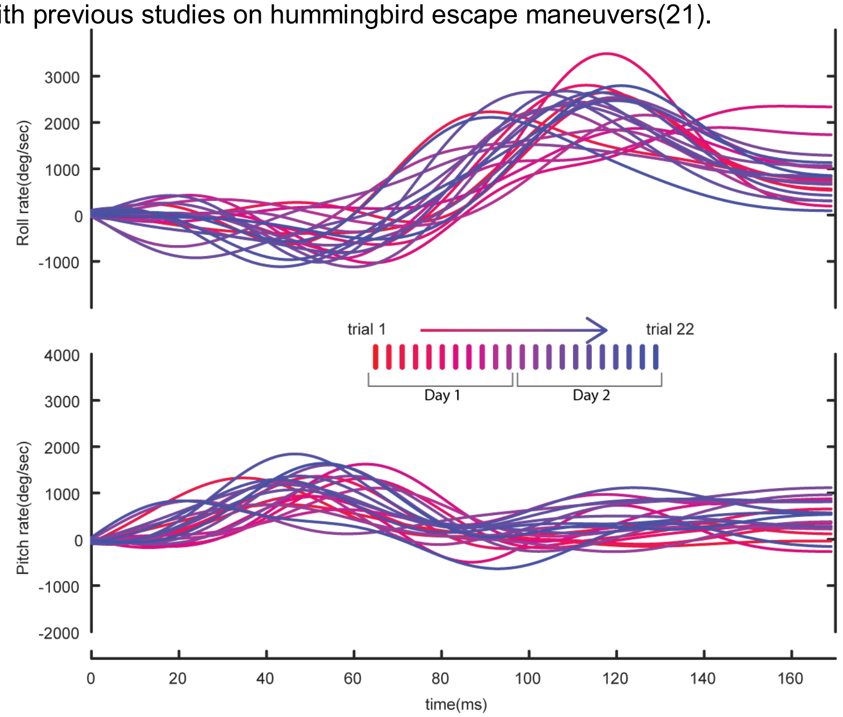


**Figure S7: No observable variation of body angular rate of hummingbird escape maneuvers over trials**. The roll and pitch rates showed no noticeable change over time, indicating consistent maneuver performance throughout the experimental trials. The lack of observable trends shows that adaptation and learning effects were not major factors in the escape responses of the hummingbirds.

**The decision either to pitch up to hover ($\mathcal{P}u\text{-}\mathcal{H}$) or pitch down to forward flight ($\mathcal{P}d\text{-}\mathcal{F}$) after the reorientation module ($\mathcal{R}$) is based on the visual input received by the bird at the critical light removal time ($t_{LR}^{c}$)**

The demonstrations during which the light was removed were categorized according to the decision to pitch up to hover ($\mathcal{P}u\text{-}\mathcal{H}$) (i.e., terminate the escape flight, category II) or to pitch down to forward flight ($\mathcal{P}d\text{-}\mathcal{F}$) (i.e., to continue the escape flight, category III) after the reorientation module($\mathcal{R}$) (Figure 1B). We found that the light removal timing $t_{LR}$ for both categories were separated by a threshold, i.e., $t_{LR}$ was less than the threshold for Category II and greater than the threshold for Categories I&III. We defined this threshold as the critical light removal time instant ($t_{LR}^{c}$). The only condition that separated category II from categories I and III was the visible light condition at the time instance $t_{LR}^{c}$, i.e., all the demonstrations in category II had visible light absent at $t_{LR}^{c}$, while for all the demonstrations in category I&III, visible light was present at $t_{LR}^{c}$.

To reject that the decision-making was based on the visual input received by the bird at a time instant other than $t_{LR}^{c}$, let's first assume that the visual input was taken at the time $t_{LR}^{input}$ different than $t_{LR}^{c}$. If $t_{LR}^{input} < t_{LR}^{c}$, then for those demonstrations in category II with the light present at $t_{LR}^{input}$ (which was possible, see Figure 1B), the bird would decide to continue the escape, which is contradictory to the decision in Category II to terminate the escape. Hence, $t_{LR}^{input} \geq t_{LR}^{c}$. Similarly, if $t_{LR}^{input} > t_{LR}^{c}$, then for those demonstrations in categories I&III with the light absent at $t_{LR}^{input}$, the bird would decide to terminate the escape, which is contradictory to the decision in categories I&III to continue the escape. Hence, $t_{LR}^{input} \leq t_{LR}^{c}$. Combining both results, we get $t_{LR}^{input} = t_{LR}^{c}$. This means that decision to $\mathcal{P}d\text{-}\mathcal{F}$ or $\mathcal{P}u\text{-}\mathcal{H}$ after $\mathcal{R}$ is based on the visual input received at the critical light removal time $t_{LR}^{c}$.

**Rate-commanded vs non-rate-commanded maneuvers**

In a rate-commanded maneuver, the body angular rate directly responds to the magnitude of the control command, i.e., the temporal profile of the body angular rate follows those of the control command. This results in a strong correlation between the control command and the body angular rate with ideally near-zero phase lag. Alternatively, during a position-commanded or an acceleration-commanded maneuver (i.e., non-rate-commanded maneuvers), the body orientation or angular acceleration responds to the magnitude of control command respectively. This results in a strong correlation between the control command and the orientation or angular acceleration with ideally near-zero phase lag.

For hummingbirds' escape maneuvers, the cross-correlation analysis revealed that the magnitude of WMPs (which can be considered as the control command) is strongly correlated with body angular rates with zero wingbeat lag (Figure 3B), instead of with body angular acceleration. Moreover, Figure 3A shows that the magnitude of WMP$^{\mathcal{P}u\text{-}\mathcal{B}}$ was nearly phase-locked with the cycle-averaged body pitch rate (rather than pitch acceleration), and the magnitude of WMP$^{\mathcal{R}}$ was nearly phase-locked with the cycle-averaged body roll rates. Taken together, these show that the hummingbird's angular maneuvers are rate commanded by the wing motion pattern.

**Assessing the degree of feedback and feedforward control through wing motion similarity**

In the field of animal locomotion, it is well-established that animals generate rhythmic movements through a combination of feedforward and feedback control mechanisms (25–28). Feedback control, which relies on state-dependent sensory information, plays a role in modulating the motor command and introduces variability into the motor commands generated by feedforward control alone. This variability stems from various sources, including the dynamics of motion, environmental factors, and sensory encoding. In contrast, feedforward control is characterized by relatively fixed motor commands, (29–31).

Based on our hypothesis, we anticipate that an increase in the degree of feedback control, such as during flight stabilization, would result in low similarity among motor commands produced in different demonstrations, compared to motor commands generated by feedforward control with a lower degree of feedback control. Direct assessment of motor command variability would require precise in situ measurement of the variability in muscle activation signals, which is currently impractical. Hence to evaluate the degree of feedforward and feedback control, we employed the similarity of wing motion patterns as an indirect measure of motor command variability (Figure 4A-i).

The interplay between feedforward and feedback control has a direct impact on the body's state during flight. Continuous feedforward control, without feedback correction, leads to a higher variance in the body's orientation. This is evident in Figure 4A, where during the evasion maneuver ($\mathcal{P}u\text{-}\mathcal{B}$), the feedforward command results in high wing motion similarity but also an increase in the variance of body orientation. However, during the reorientation phase ($\mathcal{R}(\mathcal{FB})$), with an increased degree of feedback control, reflected by lower wing motion similarity, the variance in body orientation decreases. It is well-established that hover flight is inherently unstable (32–34) and relies on feedback control for stabilization. In line with this understanding, we observed lower similarity in wing motion patterns during hovering before the initiation of the escape maneuver, indicating the involvement of feedback control (Figure 4).

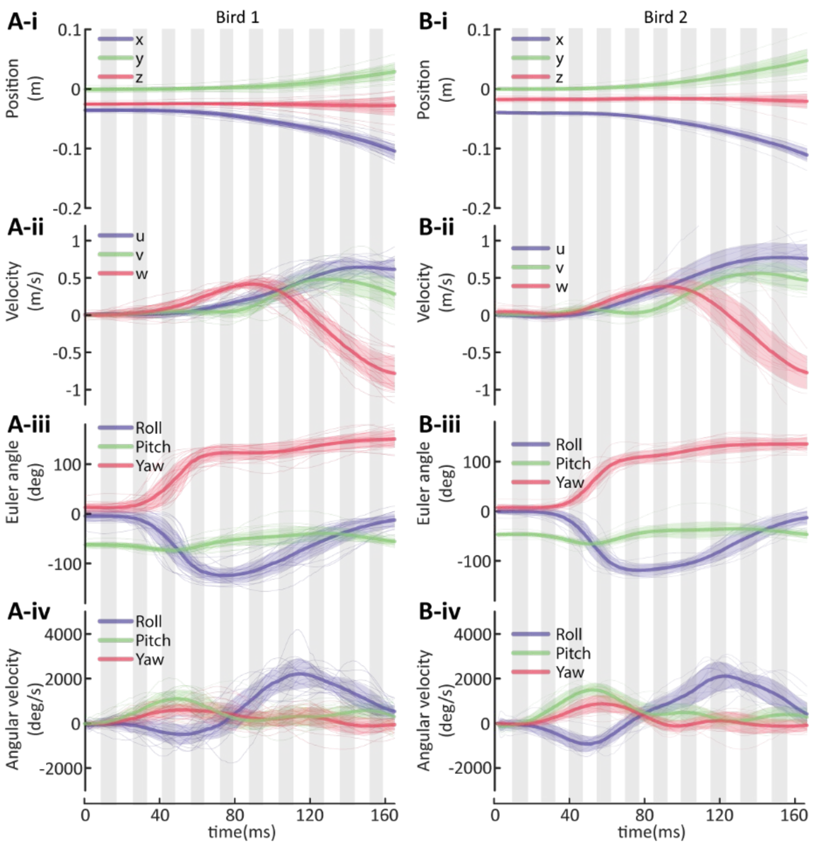


**Figure S8. Hummingbird's body kinematics during escape maneuvers.** The thick solid curves represent the averaged kinematics, thin solid curves represent the individual demonstrations, and the shaded areas indicate ±1 s.d. (22 demonstrations for bird 1, 21 demonstrations for bird 2). Vertical gray shaded areas represent downstroke.

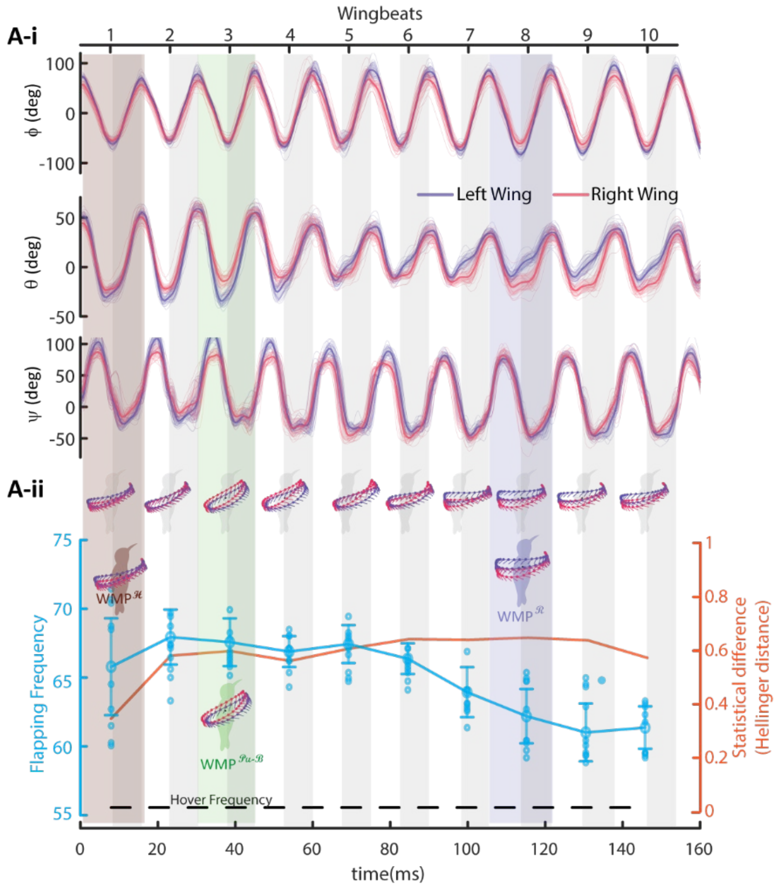


**Figure S9. Wing kinematics during escape maneuver (bird 2 data).** (B-i) Wing kinematics during $\mathscr{H} \rightarrow \mathscr{P}u\text{-}\mathscr{B} \rightarrow \mathscr{R}$ maneuvering modules. The shaded areas enclosing the curves indicate ±1 s.d. (21 demonstrations). (B-ii) Wing flapping frequency during escape flight (cyan) and individual wingbeat probabilistic distance from hovering wing kinematics (orange), which measures the kinematic difference of individual maneuvering wingbeat from averaged hovering wingbeat. The brown, green, and purple patches represent the wingbeats at which $WMP^{\mathscr{H}}$, $WMP^{\mathscr{P}u\text{-}\mathscr{B}}$, and $WMP^{\mathscr{R}}$ are defined, respectively.

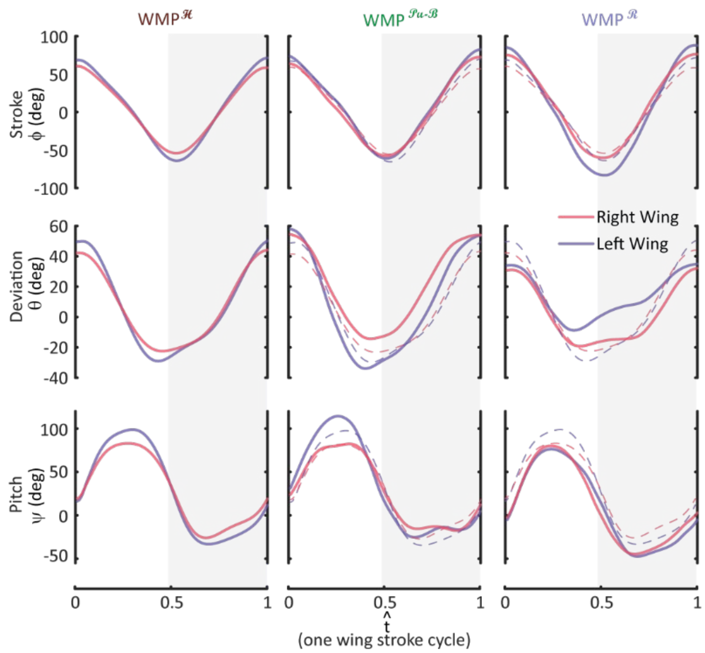


**Figure S10. Wing Motion Primitives (WMPs) during the escape maneuvers (bird 2 data).** Wing kinematic profile for WMPs, dashed curves in WMP$^{\mathcal{P}u\text{-}\mathcal{B}}$ and WMP$^{\mathcal{R}}$ represent the hovering kinematics (WMP$^{\mathcal{H}}$) for comparison. The horizontal axis represents the normalized time for a single wing stroke cycle. Downstroke periods are indicated by gray-shaded regions.

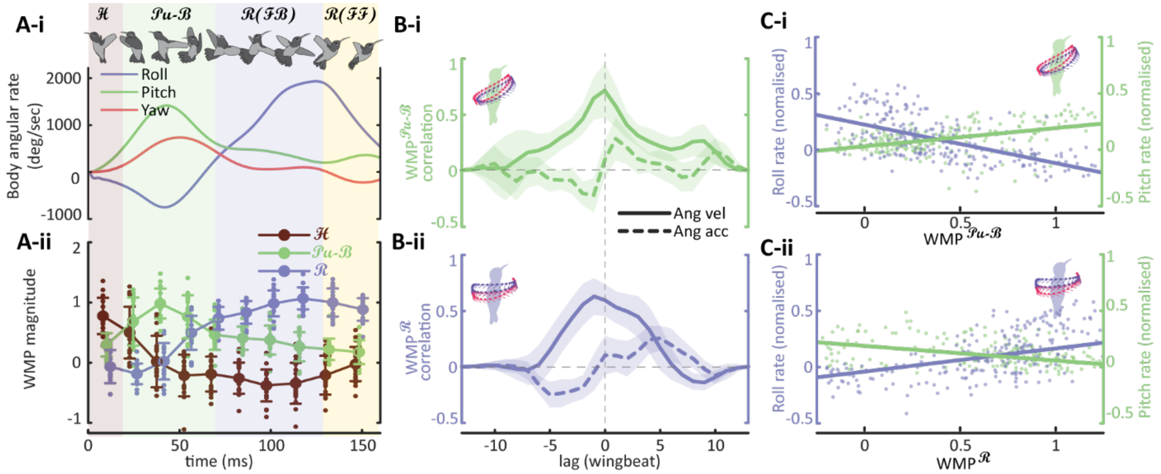


**Figure S11. Wing Motion Primitives (WMPs) are phase-locked with body angular rate (bird 2 data).** (A-i) Body angular rate in body frame. (A-ii) WMP magnitude of individual wingbeats during escape flight. The WMP magnitude follows the trend of the respective body angular rate, i.e., WMP$^{\mathcal{Pu\text{-}B}}$ follows the pitching rate while WMP$^{R}$ follows the roll rate. Background patches denote the maneuvering modules as referenced in Figure 1. (B)The cross correlation between WMPs magnitude and respective cycle-averaged angular velocity and acceleration. It is the angular velocity, instead of angular acceleration, that correlates strongly with 0 wingbeat lag and is phase-locked with WMPs. (C) Relationship between WMPs magnitude and respective normalized angular rate, and their linear regression.

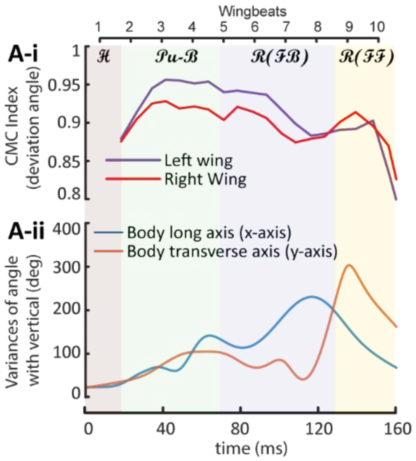


**Figure S12. Wing kinematics similarity (CMC) indicates the degree of feedforward and feedback flight control during escape maneuvers (bird 2 data).** (A-i) Coefficient of Multiple Correlation (CMC) for measuring the similarity amongst instantaneous wing kinematics of different escape demonstrations. Color background patches denote the maneuvering modules as referenced in Figure 1. (A-ii) Variances in the body orientation during escape maneuvers. The cyan curve denotes the variance in the angle between the bird's body's long axis (x-axis) and the global vertical axis, and the orange curve denotes the variance in the angle between the bird's body transverse axis (y-axis) and the global vertical axis.

**Movie S1 (separate file). Highspeed video recordings for demonstrations of category I and category III** (bird 1). In all the demonstrations visible light was present at critical light removal time instant ($\boldsymbol{t_{LR}} > \boldsymbol{t_{LR}^c}$=113 ms). Note: Category I demonstrations are the extreme cases of Category III with $\boldsymbol{t_{LR}}$=N/A.

**Movie S2 (separate file). Highspeed video recordings for demonstrations of category II** (bird 1). In all the demonstrations visible light was absent at critical time instant ($\boldsymbol{t_{LR}} < \boldsymbol{t_{LR}^c}$=113 ms).

**Movie S3 (separate file). Maneuvering modules and luminance-based responses in category I.** Visible light was present all the time. The maneuver is composed of four maneuvering modules: an initial evasive maneuver ($\boldsymbol{\mathcal{P}u}$-$\boldsymbol{\mathcal{B}}$, with body pitch up and backward acceleration), followed by a fast reorientation ($\boldsymbol{\mathcal{R}}$, with rapid roll rotation), then a dive ($\boldsymbol{\mathcal{P}d}$-$\boldsymbol{\mathcal{F}}$, with pitch down and forward acceleration,) into straight forward flight ($\boldsymbol{\mathcal{F}}$).

**Movie S4 (separate file). Maneuvering modules and luminance-based responses in category II.** The visible light was removed before the critical light removal time instant ($\boldsymbol{t_{LR}} < \boldsymbol{t_{LR}^c}$). Since visible light is absent at $\boldsymbol{t_{LR}^c}$, instead of a dive ($\boldsymbol{\mathcal{P}d}$-$\boldsymbol{\mathcal{F}}$), the bird pitched up and returned to hover ($\boldsymbol{\mathcal{P}u}$-$\boldsymbol{\mathcal{H}}$) after reorientation ($\boldsymbol{\mathcal{R}}$).

**Movie S5 (separate file). Maneuvering modules and luminance-based responses in category III.** The visible light was removed at time instant $\boldsymbol{t_{LR}}$ after the critical light removal time instant ($\boldsymbol{t_{LR}} > \boldsymbol{t_{LR}^c}$). Since visible light was present at $\boldsymbol{t_{LR}^c}$, the bird dived into forward flight ($\boldsymbol{\mathcal{P}d}$-$\boldsymbol{\mathcal{F}}$) after reorientation ($\boldsymbol{\mathcal{R}}$). But the forward flight ($\boldsymbol{\mathcal{F}}$) was terminated after a nearly fixed delay following $\boldsymbol{t_{LR}}$, and then the bird pitched up and returned to hover ($\boldsymbol{\mathcal{P}u}$-$\boldsymbol{\mathcal{H}}$) in the dark.

**Movie S6 (separate file). Wing and body kinematics of an example of demonstration from category II** (bird 1).

**Movie S7 (separate file). Summary of results.** Wing kinematics similarity (CMC) indicates the degree of feedforward and feedback flight control during escape maneuvers. Wing Motion Primitives (WMPs) magnitude are phase-locked with body angular rate showing that their angular maneuvers are rate-commanded by wing motion patterns.
